# Disentangling local adaptation and phenotypic plasticity in traits associated with altitude and temperature in widespread tropical butterflies

**DOI:** 10.1101/2025.08.29.673045

**Authors:** Tien T. T. Nguyen, Patricio A. Salazar-Carrion, Sophie H. Smith, Kimberly G. Gavilanes, Michelle Guachamin-Rosero, Gabriela Montejo-Kovacevich, Luke B. Richardson, Chris D. Jiggins, Caroline N. Bacquet, Nicola J. Nadeau

## Abstract

Climatic stratifications, in particular differences in temperature, occur along altitudinal clines. Understanding genetic and phenotypic divergence across these regions can give insight into speciation and diversification, as well as aid in our knowledge of how species may respond to possible climate change scenarios. Most past research has focused on temperate regions, yet it is in the tropics that organisms are thought to be the most vulnerable to rising temperatures. In addition, year-round stable temperatures in the tropics make altitudinal temperature variation more pronounced and increase the likelihood of local adaptation across relatively narrow gradients. Here we investigate how genetics and the environment influence a wide range of traits in two butterfly species, *Heliconius erato* and *Heliconius melpomene,* which are widespread across the neotropics and occur along the altitudinal slope of the Andes. Using ‘common garden’ rearing of over 1,000 offspring from over 70 wild females caught along an altitudinal gradient, as well as rearing of populations from either end of the altitudinal range in their reciprocal temperature environments, we find evidence of genetic, environmental, and in some cases gene-by-environment interaction effects in developmental, morphological, and thermal tolerance traits. We find parallel divergence in wing colour in both species, with wing colour darkening with increasing altitude, consistent with this playing a role in thermoregulation in these species where wing colour has mostly been linked to mimicry and mate choice. We also find evidence for gene-by-environment interactions: In *H. erato* we found local differences in heat acclimation response, with increased heat knock-out times at higher rearing temperatures found only in low altitude populations, which are exposed to the hottest temperatures. We find evidence for heritable genetic variation in most traits measured, with positive implications for adaptation to climate change, although our results suggest that selection may not act in a straightforward way on these traits.

## Introduction

Ever since Humboldt drew the attention of the scientific world to the recurrent morphologies that plants show along altitudinal gradients^1^, no matter where in the World these occur, we have considered altitude as a fundamental factor driving the diversity of form and function observed in terrestrial organisms. With consistent stratification in climatic conditions, altitude can lead to local adaptation and genetic divergence between populations at different altitudes, which in turn can result in speciation and diversification. However, despite its importance to our understanding of the origins of biological diversity across the globe, altitudinal adaptation has mostly been studied in temperate organisms^2^. Seminal work by Janzen^3^ predicted the effect of altitude on speciation and diversification to be more important in the tropics, but this has rarely been thoroughly tested^4^. In the tropics, climate not only varies rapidly with altitude but, due to the lack of seasonality, it varies little throughout the year. This creates relatively narrow and stable climatic strata^3^, which are likely responsible for the huge diversity and turnover in species composition along tropical altitudinal gradients. Despite its importance to our understanding of the evolutionary processes that lead to altitudinal adaptation and the origins and maintenance of tropical diversity, altitudinal adaptation in the topics has still attracted relatively little attention^5^.

Altitude, the vertical distance from sea-level to the top of the highest mountains, causes major and consistent changes in three key climatic features: the average ambient temperature, atmospheric pressure, and the intensity of solar radiation^6^. These changes, in turn, cause living organisms to adapt to these changing conditions. For example, many lepidopteran species show increased body size with altitude^7^ consistent with “Bergmann’s rule”^8^, which states that animals tend to have larger body sizes in colder environments, perhaps because larger body sizes increase cold tolerance^9^. Wing morphologies can also show variation with altitude. Wing loading has been found to increase across tropical moth taxa Geometridae and Arctiinae as well as in the temperate butterfly species *Coenonympha* in association with alternative flight strategies possibly due to increasing wind speeds at higher altitudes^10,11^. Wing colouration has been shown to darken at higher altitude in *Catasticta* butterfly species which was associated with increased heating rates^12^. This is consistent with a more general trend found in a range of ectotherms of darker individuals being found in colder environments, known as the thermal melanism hypothesis or Bogert’s Rule^13,14^.

Depending on the trait of interest and its ecological role, the observed altitudinal variation in a phenotypic trait can be the outcome of three modes of adaptation. These include: local (genetic) adaptation through natural selection; plasticity in developmental, physiological, morphological or behavioural responses to the environmental conditions; or a combination of both via a genotype-by-environment interaction (GxE), where the extent or direction of genetically controlled plasticity differs between populations^15^.

Evidence of local adaptation that leads to genetic and heritable differences between populations at different altitudes has been observed in many temperate butterfly species, for example *Lycaena tityrus,* when reared in ‘common garden’ (standardised) conditions, show differences in development time, heat-stress resistance, and flight duration between populations^16^. While *Colias eriphyle* exhibits gene-by-environment interactions in wing solar absorptivity across an altitudinal gradient^17^. Natural variation in trait values can often be an interplay between some degree of adaptive genetic divergence between populations coupled with some degree of adaptive phenotypic plasticity. This has been observed in butterflies such as *Melitaea cinxia* with regard to metabolic rates^18^ and in *Bicyclus anynana* with developmental growth rates^19^.

A full understanding of the nature of the phenotypic variation observed in the wild is indispensable, not only to understand the evolutionary processes of adaptation, speciation and diversification, but also in learning how living organisms may cope with long-term climatic changes (such as global warming) that are substantially altering the climate around the globe^20^. This is because while phenotypic plasticity allows for rapid adjustment to new or fluctuating environmental conditions providing immediate benefits to enhance survival, it is typically non-heritable and can be costly to maintain long term^21^. For example, a common plastic response to temperature stress is acclimation, where previous minor exposure to temperature stress leads to a priming response ^22,23^. This typically induces heat shock proteins (HSPs) more rapidly, often at a higher concentration, during future stress, thus allowing for a greater stress tolerance threshold^24^. However, HSP production is energetically costly, and constant maintenance can be detrimental to survival, as observed in *B. anynana*, where long-term exposure to heat has been shown to impair later heat tolerance^25^. Genetic variation in plasticity can be selected for and can be observed in populations across altitudinal gradients. For example, *L. tityrus* shows altitudinal differences in *hsp70* regulation in relation to rearing temperature, with lowland populations reared at higher temperatures significantly up-regulating *hsp70* levels^26^. Thus, even though the production of HSPs is energetically costly, there is a clear selection to maintain high HSP levels in warmer lowland environments rather than solely during acute stress. This highlights the importance of genetic plasticity in enabling long-term adaptation to changing environments.

Here we studied the relative roles of genetics and plasticity in determining morphological, developmental, and thermal tolerance variation across an altitudinal gradient, in two well-studied species of *Heliconius* butterflies (*H. erato* and *H. melpomene*) on the Eastern slopes of the Andes in Ecuador (Figure 1). The co-mimicking species *H. erato* and *H. melpomene* have a largely overlapping and wide distribution across the neotropics spanning an altitudinal range from sea level to just over 1000 meters above sea level (m.a.s.l.)^27^ While this altitudinal range is not particularly great, it is sufficient for populations at either end of the range to experience largely non-overlapping temperature variation, with the higher altitude sites being on average 4.1°C cooler^28^. This makes it an excellent test for the prediction that adaptation may occur over narrow altitudinal ranges in the tropics. In addition, these two species are distantly related and non-interbreeding, making them a useful study system for investigating repeated adaptation, where parallel divergence in both species can more confidently be attributed to adaptation, rather than random variation or drift^29^. Initial evidence of altitudinal adaptations within these species have thus far revealed differences in wing aspect ratio, which appears to have a genetic basis^30,31^, as well as evidence for adaptive genetic differences with altitude, some of which have been acquired via introgression from other species^29^. There is also some evidence for variation in heat tolerance, although this appeared to be primarily due to plasticity, but warrants further investigation^28^.

**Figure 1:**
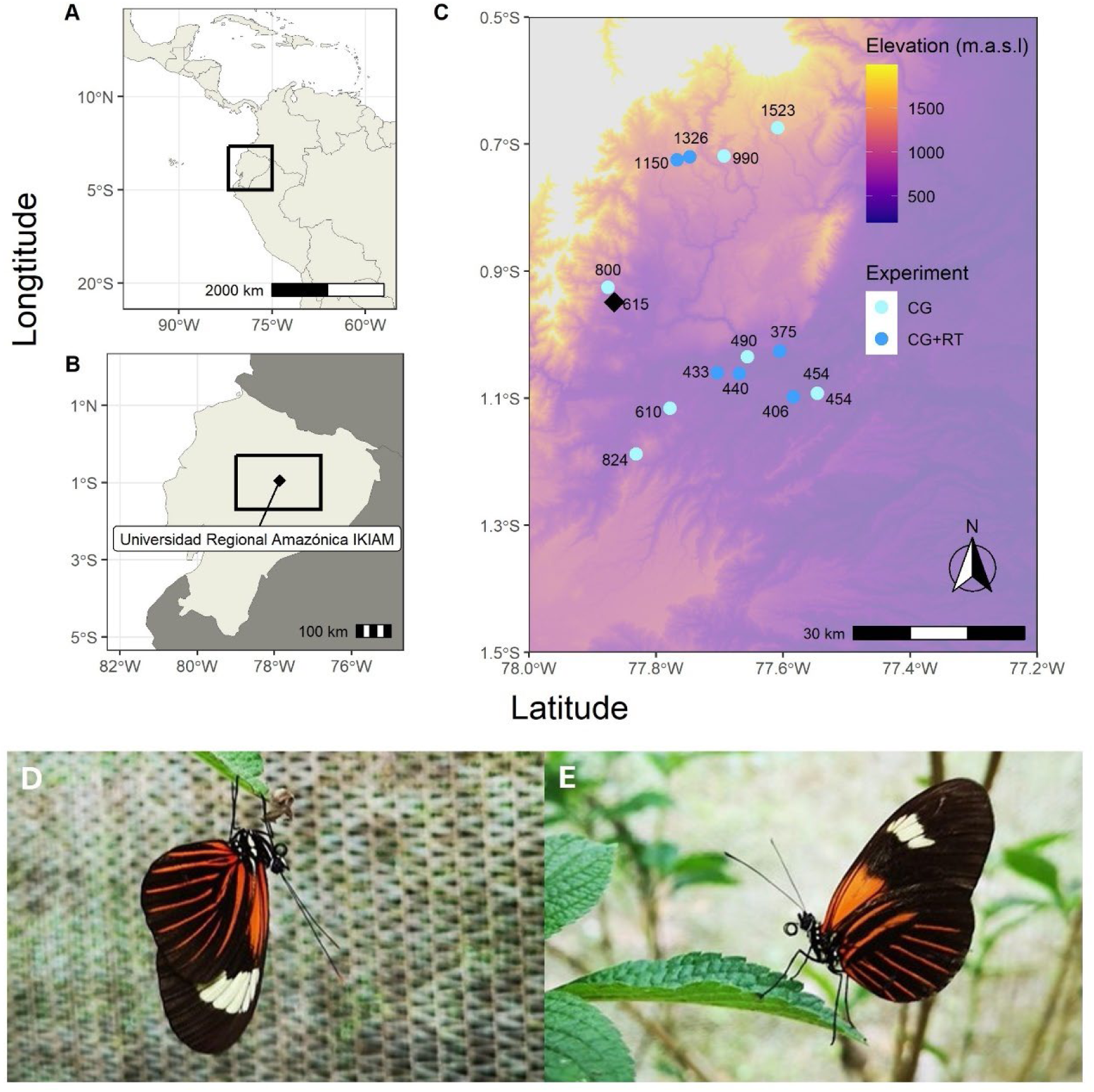
Capture sites and rearing location of butterflies sampled for the ‘Common Garden’ (CG) and ‘Reciprocal Transplant’ (RT) experiments. Location of the study within South America (A) and Ecuador (B) showing the Universidad Regional Amazonica Ikiam, where experiments were conducted, C) Locations of the rearing site (Ikiam, black diamond) and capture sites (dark and pale blue points, representing sampling sites used for both experiments and the common garden experiment only, respectively) with altitude of each point given next to it and altitude of the region shown by background colours. Sampling site altitudes ranged from 375 metres above sea level (m.a.s.l) to 1523 m.a.s.l. Photographs of the study species, *Heliconius erato* (D) and *Heliconius melpomene* (E) in the study facility at Ikiam.

Through controlled temperature rearing experiments, we aimed to disentangle the roles of genetics and plasticity in these species with regard to a range of morphological, developmental and thermal tolerance traits. The presence of genetically-controlled phenotypic differences between populations would be evidence of local adaptation and support the hypothesis that species in the tropics can and do adapt to relatively narrow altitudinal clines. In contrast, tropical species have been suggested to be less likely to show adaptive plasticity in response to temperature, due to the presence of stable year-round temperatures, meaning there should be less adaptive benefit of altering the phenotype in response to temperature^4^.

Two separate experiments were conducted to investigate the relative roles of local (genetic) adaptation and phenotypic plasticity in the two studied species. The first experiment, a Common Garden Experiment (CGE), involved individually rearing the offspring of wild-caught females, collected across a continuous altitude gradient of just over 1000 meters (Figure 1C) under common laboratory conditions that were intermediate in temperature between the collection sites at either end of the gradient (mean of 21°C). The second experiment, a Reciprocal Transplant Experiment (RTE), involved rearing offspring from wild-caught females from the two altitude extremes (Figure 1C) in growth chambers under two temperature treatments that replicated the average conditions found in the understory of these sites (means of 19°C and 24°C respectively). This was, therefore, a fully-crossed design between rearing temperature and altitude. For convenience, we will hereafter refer to butterflies reared from individuals captured below 450 m.a.s.l. as “low altitude” and those reared from individuals captured above 1000 m.a.s.l. as “high altitude”, acknowledging that this is not particularly high in the context of the Andes.

Overall, by rearing individuals from populations across a range of altitudes in a common environment, the CGE allowed us to assess the heritable effects of altitude controlling for plasticity. On the other hand, the RTE allowed us to tease apart the effects of rearing conditions (phenotypic plasticity) from heritable population differences (local genetic adaptation). In addition, due to its crossed design, the RTE allowed us to assess potential genotype-by-environment interactions (implying heritable population differences in the extent or direction of plasticity), when individuals from different altitudes showed distinct responses in the two alternative temperature treatments.

## Results

### Differences between altitudinal populations

The common garden experiment (CGE) involved rearing the offspring of wild-caught mothers in the same common temperature environment. For *H. erato*, between 29 and 45 families were included in the analyses, depending on the trait examined. Specifically, we analysed >400 individuals for development time and body size, 1549 individuals for larval survival, 802 for pupal survival, and >300 for wing morphology, colour, and thermal tolerance. For *H. melpomene*, we analysed between 20 and 25 families across traits, with >400 individuals for development time and body size, >800 for survival, and >300 for other traits.

The traits that showed substantial differences between populations across altitudes were wing colouration and heat tolerance. In both *H. erato* and *H. melpomene* the percentage of the forewing that is black increased with altitude (Figure 2A and B, Figure S1), with the best supported models for both species including altitude as an explanatory variable (*H. erato* AICcWt = 0.14; *H. melpomene* AICcWt = 0.1, Table S1). This is consistent with the thermal melanism hypothesis and findings in other species^12,14^, that higher altitude populations are often darker, possibly to improve heating ability in cooler environments.

**Figure 2:**
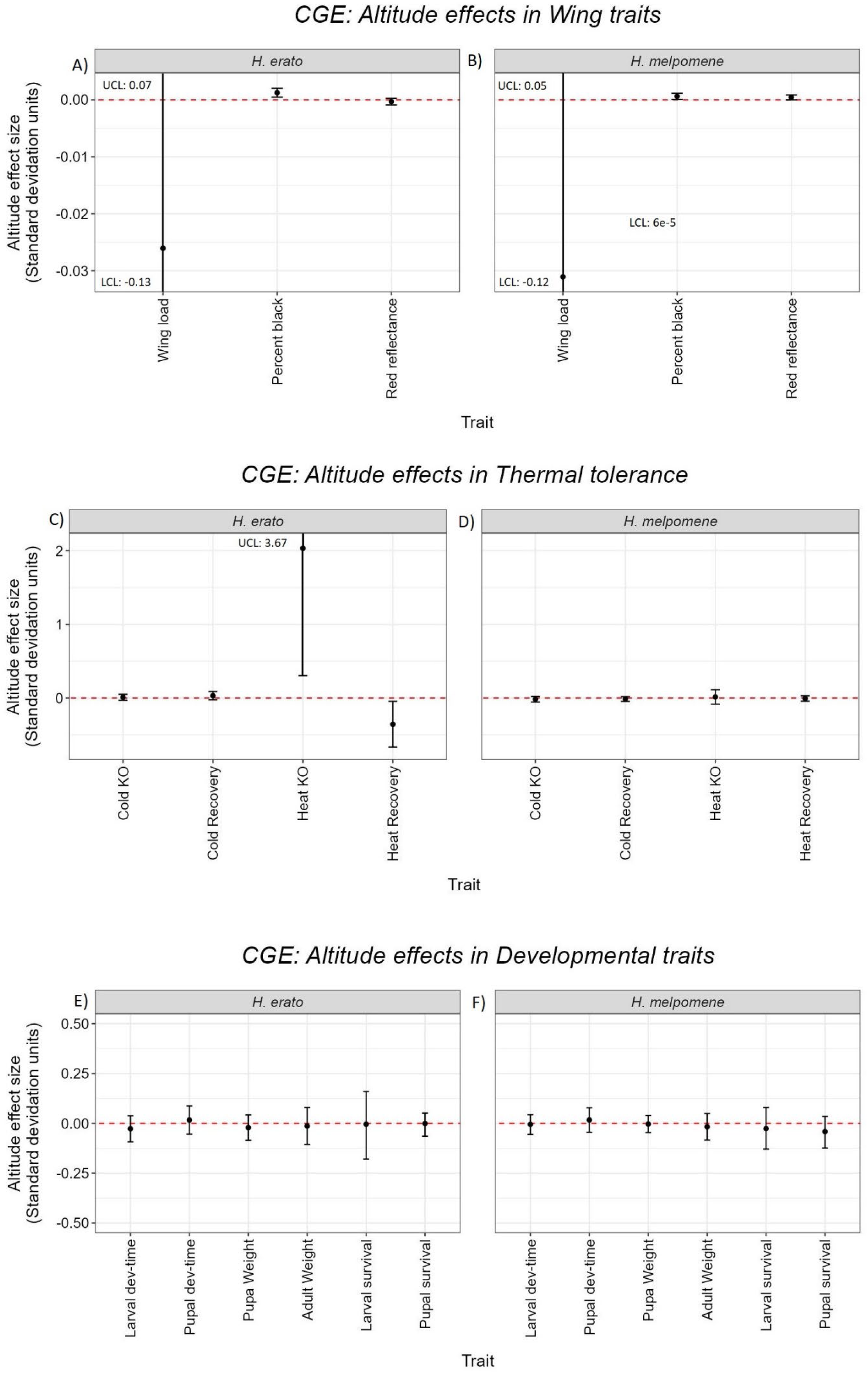
The effect of altitude on wing morphological traits in *H. erato* (A) and *H. melpomene* (B); thermal tolerance traits in *H. erato* (C) and *H. melpomene* (D); developmental traits in *H. erato* (E) and *H. melpomene* (F) calculated from the CGE.

We also measured overall reflectance (summed across all colour channels from standard photographs) of the red wing patch to determine whether it varied with altitude. In *H. erato*, there was no substantial effect of altitude on red wing patch reflectance (Figure 2A), with the highest ranked model containing only sex as a relevant explanatory factor (AICcWt = 0.34). However, in *H. melpomene*, the highest ranked model included altitude, but only when interacting with sex (AICcWt = 0.75, Table S1, Figure 2B). *H. melpomene* females showed more reflectance with increasing altitudes, while males remained relatively unchanged across altitudes (Figure S2). Overall, both species exhibited significant sexual dimorphism in the brightness of the red patch. However, only *H. melpomene* females showed variation with altitude, being paler at higher elevations, which was contrary to our expectations and inconsistent with the thermal melanism hypothesis.

For heat tolerance, *H. erato* showed differences between populations across altitudes, where individuals from higher altitude took longer to enter heat coma at 39°C (hereafter referred to as heat knock-out or KO), and less time to recover from KO when removed from the heat. Both patterns suggest an increased heat tolerance at higher altitudes, contrary to our expectations (Figure 2C, Figure S3). The highest ranked models for both traits included altitude as an explanatory factor (heat KO: AICcWT = 0.14; heat recovery AICcWT = 0.09, Table S1). For heat KO, the best supported model also contained black percentage, body weight, and their interactions with altitude. A higher black percentage increased the time to KO (effect size = 229.58x ± 204.81) while decreasing recovery time (effect size = -11.41x ± 7.41), both suggesting increased heat tolerance, again contrary to expectations under the thermal melanism hypothesis. Increased body size reduced heat recovery time (effect size = -7.70x ± 5.28), suggesting larger individuals may be more heat tolerant. However, body size also had a negative, but non-significant effect on KO time (effect size = -2.02x ± 2.08).

In *H. melpomene,* the best supported models included an effect of altitude on heat KO and recovery (Table S1), but the effect sizes were small, and the confidence intervals crossed zero (Figure 2D), suggesting the effect is not statistically significant. The models also indicated a significant effect of body size: larger individuals exhibited a shorter time to KO (effect size = – 6.73x ± 4.11), suggesting reduced heat tolerance. However, body size had no detectable effect on recovery time (effect size = 0.69 ± 2.40).

In our experiments we excluded individuals that did not knock out after 2 hours. To ensure that this did not bias our results, we repeated our analysis including these individuals with KO time set to 128 minutes, which added 70 individuals for *H. erato* and 36 for *H. melpomene*. The best supported models were largely the same (still contained altitude as an explanatory factor, Table S2) and the effect of altitude in *H. erato* was similar (effect size = 2.41 ± 1.55) and was still negligible in *H. melpomene* (effect size = 0.01 ± 0.05).

For cold tolerance, the models for both species also supported an effect of altitude on time to enter chill-coma (cold KO), and for *H. melpomene* on the time to recover from chill-coma (Table S1). However, the effect size estimates were small and crossed zero in all cases (Figure 2C and D). The models also supported an effect of body size on cold KO in both species, with larger individuals taking longer to KO, suggesting they are more cold-tolerant (*H. erato*: effect size = 3.11 ± 1.54; *H. melpomene*: effect size = 2.25 ± 1.82). This was also supported in *H. melpomene* cold recovery time, where larger individuals recovered faster (effect size = -4.72 ± 1.83).

All altitudinally-associated traits had high and significant heritability, based on the intraclass correlation coefficient (ICC) of the CGE broods (Figure 3). All individuals were reared individually under identical conditions, so similarity within broods can only be attributed to shared genetic variation or maternal effects. Therefore, heritability can be approximated as two times the ICC^32^, bearing in mind that this may be inflated by any maternal effects, which we could not distinguish in our experimental design. The heritability of black wing percentage in *H. erato* was 0.99 (ICC = 0.5, p = 1.26E-32, Figure 3A) and in *H. melpomene* 0.71 (ICC = 0.36, p = 3.57E-22, Figure 3B). For heat tolerance in *H. erato*, heritability was 0.61 (ICC = 0.31, p = 3.90E-05), and 0.51 (ICC = 0.26, p = 1.67E-03) for heat knock-out and recovery respectively (Figure 3C). Red reflectance was also significantly heritable in both *H. erato* (ICC = 0.26, p = 1.67E-14, Figure 3A) and *H. melpomene* (ICC = 0.13, p = 6.58E-08, Figure 3B). This confirms that our CGE was detecting genetic differences between populations, consistent with local adaptation to altitude. In addition to maternal effects, the estimates also assume that all individuals in each brood are full siblings. If our wild-caught females were multiply-mated, which based on knowledge of *Heliconius* mating systems is unlikely but possible^33^, this would cause an underestimation of heritability.

**Figure 3:**
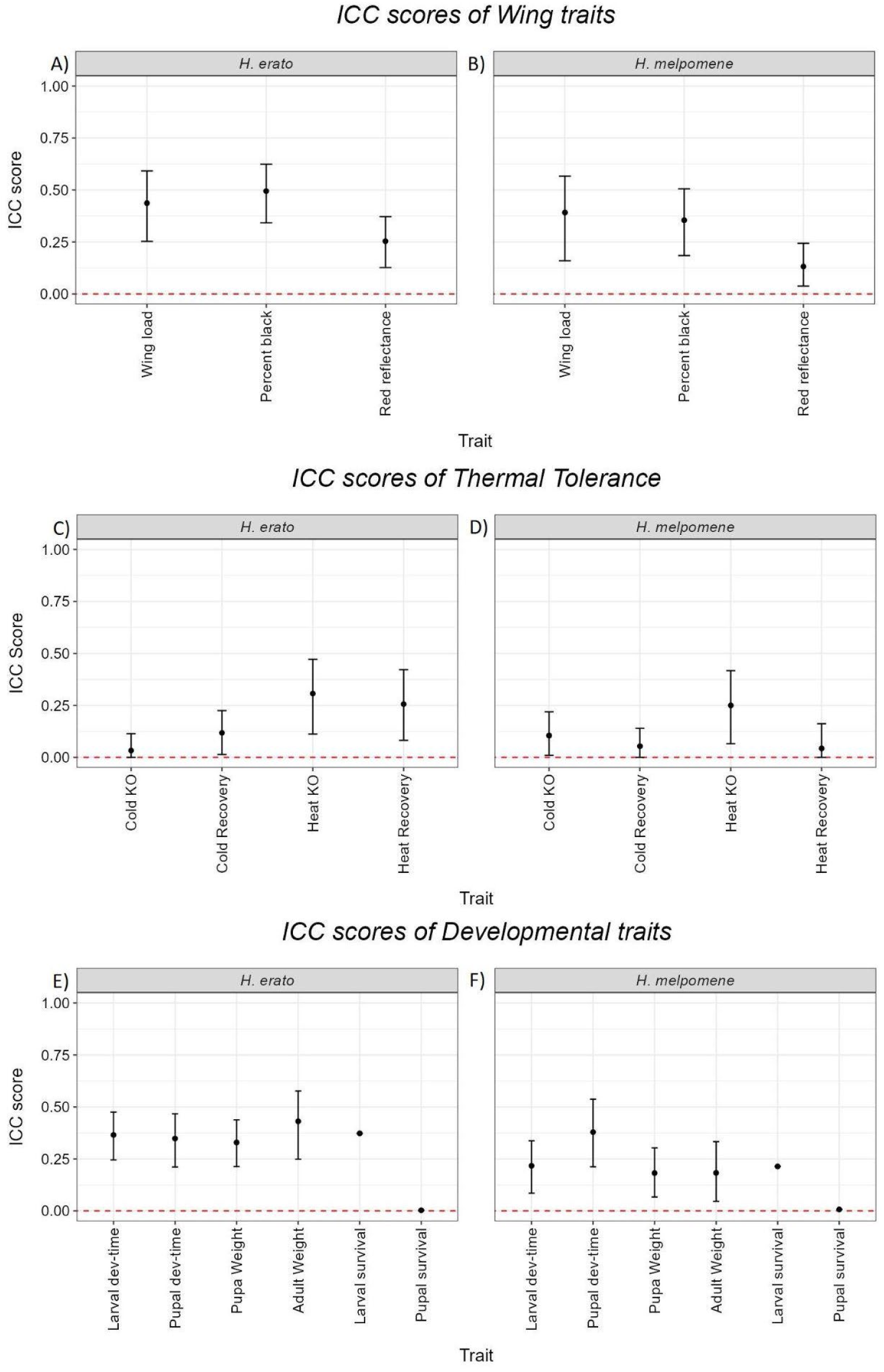
ICC scores with 95% confidence intervals of of wing morphological traits in *H. erato* (A) and *H. melpomene* (B); of thermal tolerance traits in *H. erato* (C) and *H. melpomene* (D); and of developmental traits in *H. erato* (E) and *H. melpomene* (F)

Even for traits that did not differ between populations at different altitudes, most were significantly heritable (Figure 3 C-F). Of the traits we measured, the only ones that did not show significant heritability were cold KO time in *H. erato* (Figure 3C), heat and cold recovery in *H. melpomene* (Figure 3D), and pupal survival in either species (Figure 3E and F). There were differences in larval survival of offspring coming from different mothers (*H. erato*: ICC = 0.345, p = 3.15E-72, *H. melpomene*: ICC = 0.202 p = 2.23E-13, Figure 3E and F), but there was not a significant difference with altitude (Figure 2E and F), demonstrating that any phenotypic differences we observed with altitude were not due to selective filtering of genotypes during the experiment.

### Temperature effects on traits and differences between populations

In the reciprocal transplant experiment (RTE), we reared the offspring of wild caught females from two altitudinal extremes in reciprocal temperature environments, to test for plastic effects of temperature on the same range of traits, and to determine whether any traits showed evidence of gene-by-environment interactions, whereby individuals from different populations showed different plastic responses to temperature. There were >14 families analysed in *H. erato* with >200 individuals analysed for survival and >100 for all other traits between both altitudes. In *H. melpomene*, there were >6 families analysed with >100 individuals for survival, >30 individuals for thermal tolerance and >50 individuals for all other traits. The much smaller sample size in *H. melpomene*, especially in lowland broods (Table S2), meant we had less power to discern population differences or their interaction with temperature compared to *H. erato*. See Table S2 for full details of models, and the sample sizes included for each trait.

As expected, rearing temperature had a significant effect on development time and body size in both species (Figure 4A and B, Table S3). In *H. erato* and *H. melpomene* larval and pupal development time both roughly halved at the higher rearing temperature of 24°C compared to at 19°C (effect sizes between 0.51 and 0.69, Figure 4A and B). Some marginal population differences were observed in *H. erato*, but these were within 2 AICc units of the models with temperature alone (Table S3), and also went in opposite directions in larvae and pupae, with low altitude larvae developing slightly faster under both temperatures (something that was not observed in the CGE), and high altitude pupae developing faster, but only at 19°C (Figure S4A). In *H. melpomene* there was no effect of altitude on pupal development time and the model with temperature alone was the highest ranked (AICcWt = 0.46) (Figure 4B, Table S3).

**Figure 4:**
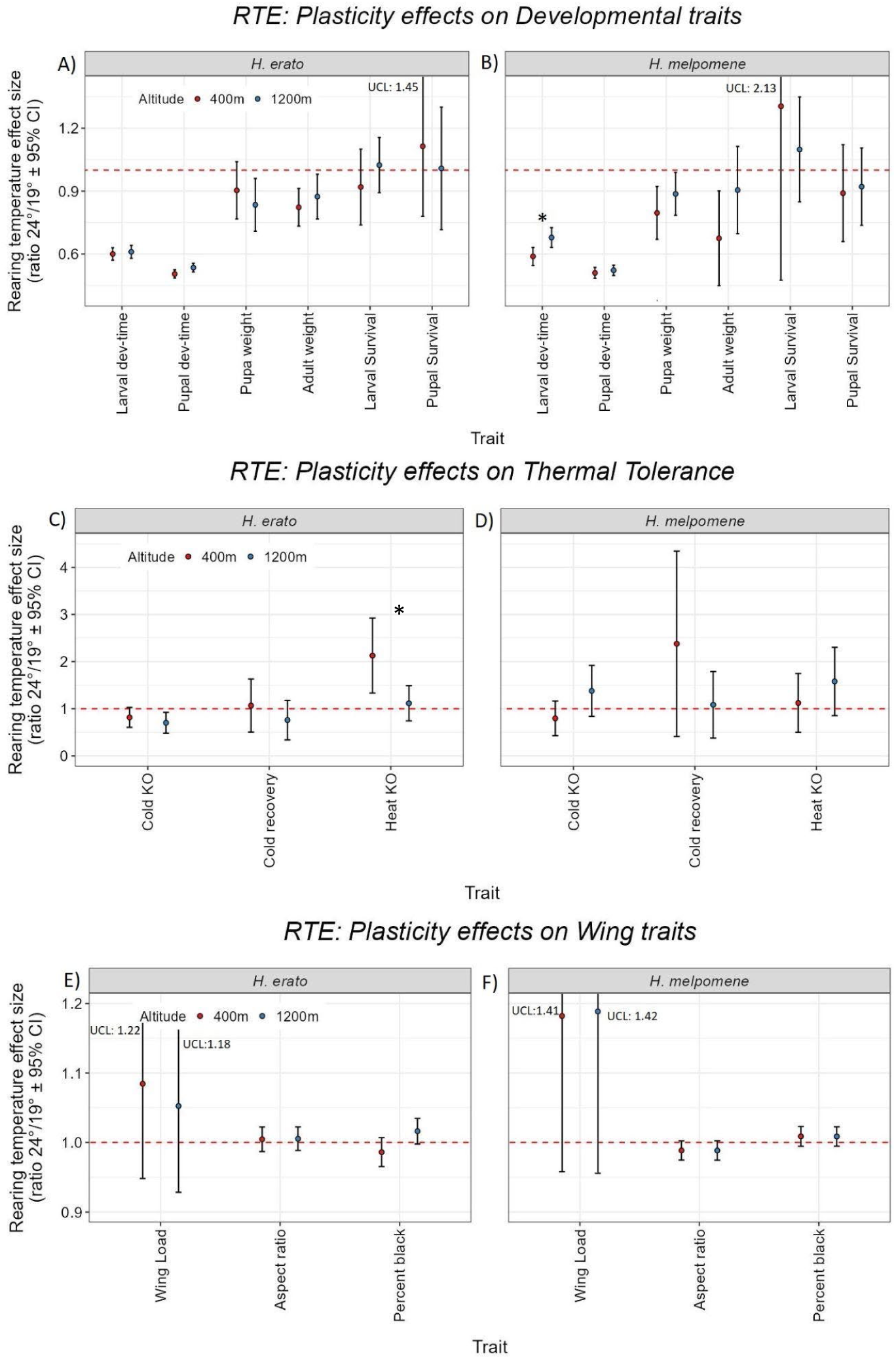
The plastic effects of increased rearing temperature (as a ratio of 24°C/19°C) in highland populations (blue) and lowland populations (red) on developmental traits of *H. erato* (A) and *H melpomene* (B) from the RTE; thermal tolerance traits of *H. erato* (C) and *H melpomene* (D); and on thermal tolerance traits of *H. erato* (E) and *H melpomene* (F). Points show the mean effect size, with error bars showing the 95% confidence intervals, and asterisk denoting substantial interaction between temperature and altitudinal population. Upper confidence limits (UCL) are given for traits where this extends off the y-axis.

However, for larval development time, the highest ranked model supported an interaction between altitude and temperature (AICcWt =0.78), with lowland populations showing greater differences in their developmental time with temperature (24°C:19°C, effect size = 0.59x ± 0.04) as compared to highland populations (24°C:19°C, effect size = 0.69x ± 0.05) (Figure 4C, Figure S5). This is consistent with an adaptive difference in *H. melpomene*, whereby high-altitude populations are able to develop faster at colder temperatures (Figure S4B, Figure S5).

We measured the effect of temperature on body size by weighing pupae one day after pupation and by weighing adults one day after eclosion, before feeding. Across both species, higher rearing temperatures tended to result in smaller pupae and adults (Figure 4A and B). There were some apparent differences in the effect sizes between populations, but the best supported model for *H. erato* pupal weight contained only temperature interacting with development time as explanatory factors (AICcWt = 0.46) and for adult weight contained temperature alone (AICcWt = 0.11, ΔAICc = 1.61). For *H. melpomene* pupa weight, the highest ranked model contained both temperature and altitude as explanatory variables (without an interaction, AICcWt = 0.56, Table S2). However, when examining effect sizes and confidence intervals there only appeared to be a temperature effect, where in both populations, pupae were smaller when reared at higher temperatures, (24°C:19°C: lowland effect size = 0.80x ± 0.13, highland effect size =0.887x ± 0.102, (Figure 4B)) with no clear altitudinal effects (Table S3). For adult weight in *H. melpomene*, our highest ranked model again contained an interaction between temperature and altitude (AICcWt = 0.27) but this interaction effect was marginal as this model was not stronger than the model without the interaction term (Delta_AICc = 0.48, AICcWt = 0.21). Therefore, the results for *H. melpomene* are similar to *H. erato*, with a lack of strong evidence for population differences in the effects of rearing temperature on body size.

We found evidence of cold acclimation *H. erato,* but not *H. melpomene* (Figure 4C and D), whereby individuals reared at cooler temperatures took longer to enter chill coma than those reared at warmer temperatures (Figure 4C). This acclimation was only observed in highland populations (24°C:19°C: 0.70x ±0.22), and not the lowland populations (24°C:19°C = 0.81x ±0.21) (Figure 4C). However, the interaction was not significant as the highest ranked model had temperature as the sole explanatory factor in time to chill coma (AICcWT = 0.51). We also found evidence of heat acclimation in *H. erato*, but again not in *H. melpomene*, whereby individuals reared at warmer temperatures took longer to enter heat coma than those reared at lower temperatures (Figure 4C and D). This effect was only seen in the low altitude populations (24°C:19°C: 2.13x ± 0.79) not those from higher altitude (24°C:19°C: 1.12x ± 0.37) (Figures S6, S7), and this population difference was significant, with the highest ranked model including the interaction between temperature and altitude (AICcWT = 0.67, Table S3). This is consistent with adaptive differences in plasticity between populations, with lower altitude populations being better adapted to respond to higher temperatures.

In *H. erato* there was also evidence for an effect of rearing temperature on the percentage of the wing that was black, in addition to the population difference that was found in the CGE. The population difference was only found when individuals were reared at 24°C, being darker in the higher altitude population (400m:1300m = 0.95x ± 0.03) (Figure S8A), but there was no difference in black wing percentage between populations when reared at 19°C (400m:1300m = 0.98x ± 0.03) (Figure S8A). This was due to greater plasticity in the higher altitude population, with butterflies being darker when reared at higher temperatures, which was not found in the low altitude population (Figure S8A). Our highest ranked model supported this interaction (AICcWt =0.19) and also suggested an interaction effect between temperature and sex, where females had greater black percentage at higher rearing temperatures in comparison to males. Therefore, the population difference observed in the CGE is complicated by differences in the extent of plasticity in wing colour in response to rearing temperature. In *H. melpomene* we did not detect any effect of rearing temperature on percentage black of the wing. Red reflectance was not measured in the RTE due to mechanical damage in the red patch.

We again did not find any difference between populations in larval or pupal survival (Figure S4), demonstrating that selection operating during our experiment was not a factor in explaining between-population differences (or lack thereof). We also did not find any difference in survival between populations at the different rearing temperatures (Figure 4A and B), suggesting that the mean temperatures at the different altitudes are not causing differential selection between populations during the larval or pupal stages.

## Discussion

Wing colour in *Heliconius* has been extensively studied, primarily in the context of Müllerian mimicry, and several genes that underlie pattern variation have been identified^34,35^. Here we have found evidence in both species of quantitative variation in the proportion of black pigmentation on the wing associated with altitude, evidence that environmental factors, in addition to predation, influence selection on wing colour in these species. These traits were also highly heritable, raising the interesting possibility that they may be controlled by some of the major colour pattern controlling genes that are known in these species, as has been found for other quantitative colour pattern variation^36^. Similar trends of darker colours with increasing altitude have been found both within and between species in many insects^37^ and are generally thought to be explained by thermal melanism, whereby darker individuals warm up faster when exposed to radiative heat^14^. Most of this data has come from temperate systems, where cold temperatures are likely to be more extreme and limiting, so it is interesting that we also observe this trend in tropical insects. A previous study of tropical butterflies in Ecuador found conflicting results, with one genus (*Catasticta*) having darker species at higher altitudes, while another (*Leptophobia*) showed the opposite trend^12^. We are not aware of any previous studies that have found a similar within-species trend in tropical butterflies.

While the colour variation we observe is consistent with thermal melanism, in *H. erato* this pigmentation had the opposite effect to expectations in the thermal tolerance experiments, where it appeared to increase heat tolerance. One explanation for this could be the artificial conditions of our experiment, where heating was purely via conduction and convection, while in nature, solar radiation is a major heat source^38^. In temperate systems, cold temperature is likely to be a strong selective agent at higher altitudes, but in tropical systems increasing solar radiation may be more important. Increasing solar radiation has been suggested to explain why some species may show reversed trends of lighter colour with higher altitude, to prevent overheating under high solar radiation^39^.

In both species we find interesting sex differences in the reflectance of the red wing patch, consistent with previous work on these species showing subtle sexual dimorphism^40^, which may function in mate choice. In *H. melpomene* we also find an interaction between sex and altitude, with *H. melpomene* females from higher altitude being more reflective. This would be consistent with females adapting to increased solar radiation. Males may be more constrained by sexual selection, or less adapted to local conditions due to their higher dispersal^41^.

Overall, we find heritable differences between populations in wing colour, building on our previous working demonstrating heritable differences in wing shape between the same populations^30^. While the differences in wing colour were clear in our common garden experiment, when reared under different temperatures in our reciprocal transplant experiment, we found that *H. erato* showed thermally plastic differences in wing colour that differed with population and sex, which could make it hard to detect the colour difference in the wild, highlighting the benefits of disentangling genetic and environmental effects under controlled conditions.

A key assumption of Janzen’s hypothesis^3^ that altitude has a greater impact in the topics, is that tropical species will show narrow, altitudinally stratified thermal tolerance ranges^4^. We found limited support for this, with no difference in thermal tolerance between populations in *H. melpomene*. We did find significant differences between populations in heat tolerance in *H. erato*. However, in our common garden experiment these differences were in the opposite to expected direction (increased thermal tolerance in the high-altitude populations) and the opposite direction to that found previously by Montejo-Kovacevich et al.^28^ in wild-caught individuals. This is likely due to the plasticity in this trait that we detected in our reciprocal transplant experiment, which also revealed differences in the extent of plasticity between populations. The low-altitude populations show an acclimation response, whereby rearing at higher temperatures causes them to be more heat tolerant as adults, but this response is not seen in the high-altitude populations. Therefore, when reared at 24°C, we find the expected population difference, with low-altitude populations being more heat tolerant. The opposite trend that we find in the common garden experiment is therefore likely due to low altitude populations becoming less heat tolerant when reared at the comparatively cool temperature of 21°C, while the high-altitude population remains relatively heat tolerant. This may also explain why Montejo-Kovacevich et al.^28^ found no difference between populations in their common-garden experiment, as these were conducted outdoors at warmer and more variable temperatures.

Lack of a heat acclimation response in the high-altitude populations could be due to greater daily variation in temperatures seen in the high-altitude tropics, due to higher solar radiation^4^, making it less beneficial for high-altitude populations to respond plastically to developmental temperature, while at lower altitudes, wet and dry season temperature variation may be sufficient to make long-term acclimation beneficial. This goes against expectations that tropical species should not show acclimation responses due to the relatively stable temperature environments they inhabit^4^, although this was supported in *H. melpomene* where we did not detect any differences in heat knock-down with rearing temperature.

Interestingly we also found evidence of cold acclimation *H. erato*, where individuals reared at cooler temperatures took longer to enter chill coma than those reared at warmer temperatures. This effect has been commonly observed across insect species including some tropical species like *Drosophila melanogaster*^42^ and the butterfly *Bicyclus anynana*^43^. It is particularly interesting here, given that the cold temperatures used in our knock-out experiments are below anything these butterflies would experience at their capture locations^28^. The distribution of *H. erato* extends beyond the tropics into regions where they could experience extreme cold^27^. Thus, it is possible that there is some gene flow from these populations that maintains this plastic response. This could be somewhat similar to the situation in stickleback fish, where genetic variation shared across their broad geographical range has facilitated repeated adaptation to new environments^44^. We did not observe any cold acclimation effect in *H. melpomene*, consistent with their differing latitudinal ranges.

In temperate butterflies, developmental rates often increase with altitude^45^ to compensate for shorter seasons and cooler temperatures^2^, although due to differences in voltinism, some species show the opposite trend, with slowed development and one generation per year at higher altitude, and faster development to allow for more generations at lower altitude^16^. We did not observe any difference in developmental rate between populations in our common garden experiment, suggesting that in tropical systems the lack of seasonality may relax selection on development time, as life-cycles do not need to be completed within a fixed time-window. Nevertheless, *H. melpomene* did show faster larval development in high compared to low altitude populations when reared at 19°C, suggesting that selection has favoured the ability to develop faster at cooler high-altitude temperatures in this species. Faster developmental times may be selected to reduce vulnerability during juvenile stages to predation as well as allowing for earlier dispersal as adults, which could be beneficial when competing for mates and hostplants^46^. A possible explanation for the difference between species it that *H. melpomene* is a host-plant specialist, which may mean that they experience greater resource competition than *H. erato*, which is more generalist.

Many studies report differences in body size with altitude in insects^2,31,47^, often following Bergmann’s rule^8^ where warmer environments contain individuals or species with smaller body size. This has often been suggested to be adaptive based on larger individuals being more thermally buffered by greater surface-area to volume ratios^48,49^. While this is logical for endotherms, the predictions in ectotherms are less clear and trends found in the wild are likely to be highly biased by differences in growth-rates^47,49,50^. We did not detect any difference in body size between populations when reared under common-garden conditions, suggesting that previously reported differences in size with altitude in these species are due to plasticity^31^. Indeed, we found a strong effect of rearing temperature on body size that followed the temperature size rule, a common reaction norm where increases in rearing temperature decrease body size^51^. We found this was tightly coupled with development time, where shorter development times (caused by warmer temperatures) also led to smaller pupae. Whether this plasticity provides an adaptive advantage or is just a general biological consequence^52^ has been contested^53^. The effect of body size on thermoregulation in insects appears to be highly variable^54,55^. Our results were generally consistent with larger-body sizes being beneficial in cooler temperatures, and so with the observed plasticity being adaptive, as in both species larger individuals were more cold-tolerant and in *H. melpomene* larger individuals were more susceptible to heat, although in *H. erato* there was some evidence that larger individuals were more heat tolerant.

Consistent with Janzen’s hypothesis^3^ we do find evidence for adaptive differences between populations of two species of tropical butterflies across a relatively narrow altitude range (∼400 m.a.s.l. to ∼1200 m.a.s.l.). However, these differences are primarily in morphological traits, principally wing colour and shape. Further work is needed to understand exactly how these traits confer adaptive benefits between populations, but their high heritability and parallel divergence between these two species, suggests they are likely to be adaptive.

In contrast we find little evidence for fixed differences in physiological and developmental traits between populations, although we do find high heritability for these traits. Lack of divergence in traits with high heritability would suggest that selection is absent or too weak to overcome gene flow across the altitudinal cline^56^. Alternatively, selection may be acting in a more complex way on the levels of plasticity in these traits. In support of the latter, we find intriguing genotype-by-environment interactions in thermal tolerance and developmental rate in *H. erato* and *H. melpomene* respectively, highlighting the benefits of reciprocal-rearing experimental designs for teasing apart genetic and environmental effects on traits. The very different responses we find between species in these traits could reflect differences in ecological or evolutionary constraints between the two species and may influence how each is able to respond to changing environments. The variation in heat acclimation that we observe in *H. erato* may be particularly beneficial in helping this species to respond to climate change, while the lack any acclimation response in *H. melpomene* may make it more vulnerable. Nevertheless, the presence of genetic variation for thermal tolerance in both species, suggests that they have the potential to adapt to changing environments and climate change.

The presence of altitude-associated phenotypic differences is consistent with previous work in these same populations, identifying signatures of adaptive genomic differences^29^. The next step will be to understand how the genetic variation controls the phenotypic variation we have found. This will allow for a deeper understanding of the adaptive potential of these traits and how they may be influenced by current and future climates.

## Methods

### Experimental design

The Common Garden Experiment (CGE) involved rearing the offspring of wild-caught females from both species under consistent, controlled conditions. These females were collected across a continuous altitude gradient ranging from 380m to 1250m above sea level for *H. erato* and 380m to 1500m for *H. melpomene* (Figure 1C). Rearing took place under controlled laboratory conditions at the Amazonian Regional University Ikiam (Figure 1B). Laboratory temperature was set to be approximately intermediate between the temperatures observed at either end of the altitudinal range^28^: 21.2 ± 1.1 °C (mean ± sd). Humidity was monitored but not strictly controlled, and the light regime followed the natural day-night cycles at this latitude (∼12:12 hours).

The second experiment, a reciprocal transplant experiment (RTE), involved rearing offspring from wild-caught females from two altitude extremes, approximately 400 ± 50m (lowland) and 1200 ± 100m (highland) above sea level (Figure 1C). Offspring from both altitudes were reared in growth chambers under two temperature treatments that replicated natural highland and lowland forest understory conditions^28^: 19.1 ± 1.7 °C for ∼1200m and 23.5 ± 2.1 °C for ∼400m. A 12:12 hours day-night regime was implemented in both growth chambers. Chamber temperature conditions closely replicated natural conditions (Figure S10)

Collection sites for the mothers were restricted to the geographical range of the co-mimetic subspecies *H. erato lativitta* and *H. melpomene malleti*, to avoid potential confounding effects associated with subspecies colour pattern variation. Mothers were kept in shaded outdoor insectaries at Ikiam (600m above sea level). Eggs were collected daily in the late afternoon at the same time, moved to the lab, and separated into individual rearing containers. Caterpillars were provided with a cutting of a suitable food plant for each species (*Passiflora punctata* for *H. erato* and *P. edulis* for *H. melpomene*) and allowed to feed *ad libitum*. Traits associated with development, thermal tolerance, and wing morphology were measured for each reared individual.

### Developmental parameters

Eggs, larvae, and pupae were checked daily throughout the experiments to accurately record mortality, and developmental time from hatching to moulting into 5th instar larvae, pupation and eclosion into adult butterflies. Pupa weight was measured one day after pupation, and adult weight was measured one day after eclosion, before feeding.

### Thermal tolerance

Adult offspring were tested for their tolerance to thermal extremes of hot and cold. All thermal tolerance tests were conducted in the ‘common garden’ rearing lab at 21°C, and researchers carrying out the experiment were blind to the butterfly’s altitude of origin.

For CGE adults, cold tolerance tests were conducted on the second day after eclosion and heat tolerance tests were conducted the following day. Whereas for the RTE, butterflies were randomly assigned to either the cold or heat tolerance test (not both) conducted on the second day after eclosion (as these individuals were used for a subsequent gene expression study).

Cold tolerance was tested by exposing the butterflies to 5 ± 1 °C for 10 minutes, measuring the time taken for the butterfly to exhibit chill coma response^57^ (or time to cold knockout (KO)) and then the time taken to recover, which was characterised by the ability to support itself on 3 or more legs.

Heat tolerance was tested as heat knockout (KO) time at 39°C (± 1°C) (and recovery, in common garden only) following the methods of Montejo-Kovacevich et al.^28^, with minor modification. After the heat tolerance experiment, all butterflies were killed and preserved. Butterflies that did not knockout after 120 minutes were excluded from the analysis. To ensure that this did not bias the results we also conducted an analysis where these individuals were included in the analysis with KO time set to 128 minutes.

### Wing Measurements

Detached wings were photographed dorsally with a Nikon D7000 APS-C DSLR with fixed settings of: 40 mm f/2.8 lens, f/10, 1/60, ISO 100, under standardised conditions with an X-Rite passport colorchecker and ruler included. Damage to the wings was assessed by eye, with wings showing more than 10% damage removed from our analyses for the GCE samples, but all samples were kept in RTE as long as the whole wing segment was present. Custom scripts were developed in Adobe Photoshop 2021 to automate colour correction and background removal. Wing size, and shape were measured from preserved wings analysed as in the methods in Montejo-Kovacevich et al.^31^. ‘Wing load’ was measured by dividing total adult weight (mg) by wing area (mm^2^) to give mg of weight per mm wing (mg/mm^2^).

Black wing proportion as well as red brightness were obtained using the ‘ColourDistance’ R package through a k-means clustering approach (Figure S9). We specified that the images contained three distinct colours, as both species have distinct patches of black, red and yellow. ColourDistance clusters all pixels most similar to one another into the defined number of groups and calculates the proportion and mean R.G.B. values for each group^58^. Brightness of the red region (hereafter red brightness) was calculated by combining mean red, green and blue values in the red region and dividing by three to get an estimate of percentage reflectance (within the range of the camera’s sensitivity).

All wing photographs are available on Zenodo, under the following DOIs: *Heliconius erato* dorsal wing images from common garden experiment, doi:10.5281/zenodo.16688551, doi:10.5281/zenodo.16690029, doi:10.5281/zenodo.16690550, doi:10.5281/zenodo.16691109, doi: 10.5281/zenodo.16691604, doi:10.5281/zenodo.16691924, doi:10.5281/zenodo.16692034; *Heliconius melpomene* dorsal wing images from common garden experiment, doi:10.5281/zenodo.16781404, doi:10.5281/zenodo.16781503, doi:10.5281/zenodo.16781637, doi:10.5281/zenodo.16781731, doi:10.5281/zenodo.16781817, doi:10.5281/zenodo.16781920; *Heliconius erato* dorsal wing images from reciprocal transplant experiment: doi:10.5281/zenodo.16762443, doi:10.5281/zenodo.16762500; *Heliconius melpomene* dorsal wing images from reciprocal transplant experiment, doi:10.5281/zenodo.16762538, doi:10.5281/zenodo.16762573

### Data analyses

Variation in each measured trait was modelled using Linear Mixed-Effect or Generalised Linear Mixed-Effect Modelling depending on the statistical nature of the trait (e.g., Gaussian distribution for morphological measures like body mass and Gamma distribution for ‘waiting-time’ variables such as development times).

Explanatory variables incorporated in the models varied depending on the experiment and trait analysed. In both experiments, the set of explanatory variables of interest was implemented explicitly in the experimental design. In the CGE, ‘altitude’ (modelled as a continuous variable) was the key effect of interest, while in the RTE ‘rearing temperature’ (19°C and 24°C treatments), ‘altitude’ (categorical: 400m and 1200m) and the statistical interaction between them were the effects of interest. Additional potentially explanatory variables were considered for each trait; for instance, ‘sex’ for traits measured in adults (which can be sexed), and ‘observer identity’ for traits affected by subjective observer bias (e.g., cold- and heat-knockdown times). In addition, because in both experiments a variable number of offspring of the same mother were reared, we modelled the effect of relatedness by including mother as a random effect. This random effect not only allowed us to account for the lack of statistical independence between offspring of the same mother (full sibs), but also allowed us to estimate heritability for the measured traits (see below).

For each trait, we followed the same modelling protocol. We first generated a list of nested candidate models combining all explanatory variables plus any pairwise statistical interaction considered relevant for each trait (for the RTE, the interaction between ‘rearing temperature’ and ‘altitude’ was always incorporated, since it was a key parameter of interest). We then fit the candidate models to the data, ranked them according to their AICc scores^59^, and selected the model with the lowest AICc value that would also contain the set of explanatory variables of interest in each experiment. We used this selected model to estimate the effect sizes of interest in each experiment. The model selection process was also used to assess which potentially explanatory variables showed evidence of an effect, and which did not. Explanatory variables with a likely effect on a trait were taken as those contained in the model with the lowest AICc (which we term the “top ranked model”, but not necessarily the one we selected to estimate effect sizes), or in the set of models ranked within two AICc units from the top ranked^59^. Model fitting was carried out using the *R* package *lme4* (v.1.1-31)^60^, and model selection facilitated with *AICcmodavg* (v.2.3-2,)^61^.

With the CGE data only, the random effect of ‘mother’ estimated with the selected model was used to calculate the intraclass correlation coefficient (ICC) between offspring of the same mother, following Nakagawa and Schielzeth^62^, using package *rptR* (v.0.9.22)^63^. This ICC was in turn used to estimate each trait’s heritability as 2 * ICC, measuring the upper limit of heritability, based on Falconer and Mackay^32^. Re-mating is rare in the wild in the two species studied^33^, so we assume that individuals from the same mother are full siblings.

Finally, to compare the effects of altitude on the various traits measured in the CGE, we recalculated the slopes of each trait in relation to altitude, after standardising the original trait values (mean = 0, sd = 1) and re-fitting the selected models. For visual comparisons, we plotted these standardised slopes (effect sizes in sd units) next to each other. For the RTE, in contrast, to compare the effects of rearing individuals from the same altitude in different temperatures (phenotypic plasticity), or to compare individuals from different altitudes reared in the same temperature (local genetic adaptation), we calculated the relevant effect sizes as the ratio between the estimated marginal means (as predicted by the selected models) for individuals in the corresponding temperature-by-altitude combinations. These effect sizes were estimated using the package *emmeans*^64^.

All statistical analyses were conducted in R version 4.+ (R core Team). All graphs were produced with ‘ggplot2’^65^, and maps required additional packages “rnaturalearth”^66^, “raster”^67^, “sf”^68,69^ and “ggrepel”^70^. The data and scripts used for analyses are available on Zenodo, doi: 10.5281/zenodo.16754981

## Acknowledgements

We thank the many research assistants and interns from Ecuador and the UK whose dedication was essential to this work, especially Gladis Grefa, Jimmy Velasteguí, Pamela Chamba, Karen Muñoz, Carlos Robalino, Sylvie Tranter, Michelle Campaña, Imogen Elliott, Jamal Kabir, Edwin Ochoa, Tamia Salazar, Andrei Flores, Franz Chandi, Jonathan Wood, and María José Sánchez, as well as other students from Ikiam who contributed to caterpillar rearing and data collection. Sihang Lai, Ryan Adams, and Megan Webb provided assistance with wing photography. We thank Alexandra Durán, Jorge Batres, and Carolina Proaño (Universidad Regional Amazónica Ikiam) for long-standing support with logistics and laboratory access, and the Ikiam technical staff—especially Nina Espinoza de los Monteros, Andrea Carrera, and Andrea Salgado—for broader assistance. We thank Carlos Valle-Piñuela (Ministerio del Ambiente y Agua del Ecuador) for his support in processing the framework contracts for access to genetic resources (MAE-DNB-CM-2018-0088 and MAATE-DBI-CM-2021-0176), as well as export permits, which enabled this research. This work was supported by a NERC grant (NE/R010331/1) to N.J.N. and C.J. and a NERC ACCE studentship to T.T.T.N. (2302726).

## Extended Data

**Figure S1:**
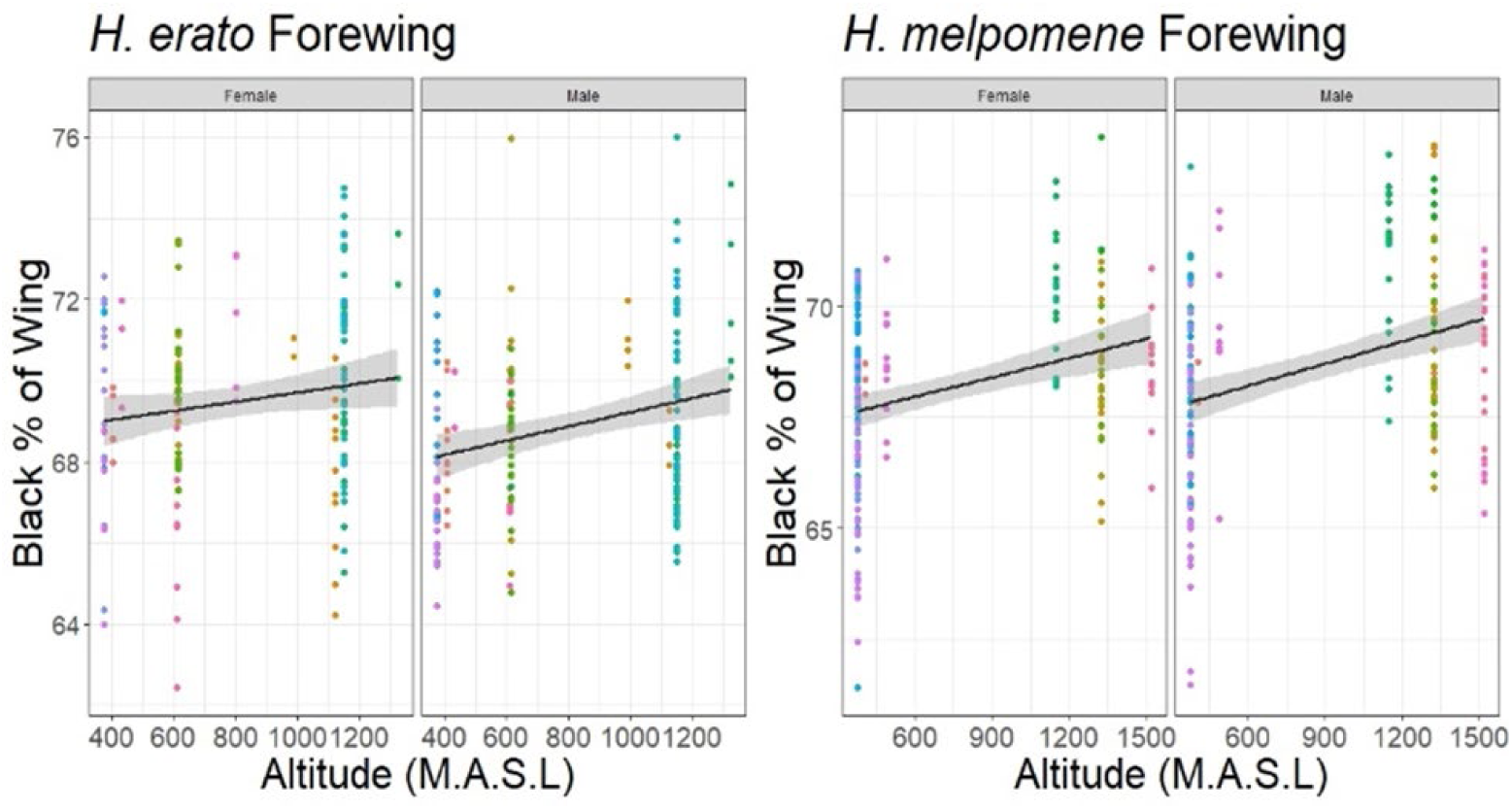
The relationship between wing black percentage of the wing and altitude of the source population in both *H. erato (*A) and *H. melpomene* (B). Points are coloured by mother ID (points of the same colour are siblings). Trend lines are based on the raw individual data (not the effect sizes from the full model of all effects).

**Figure S2:**
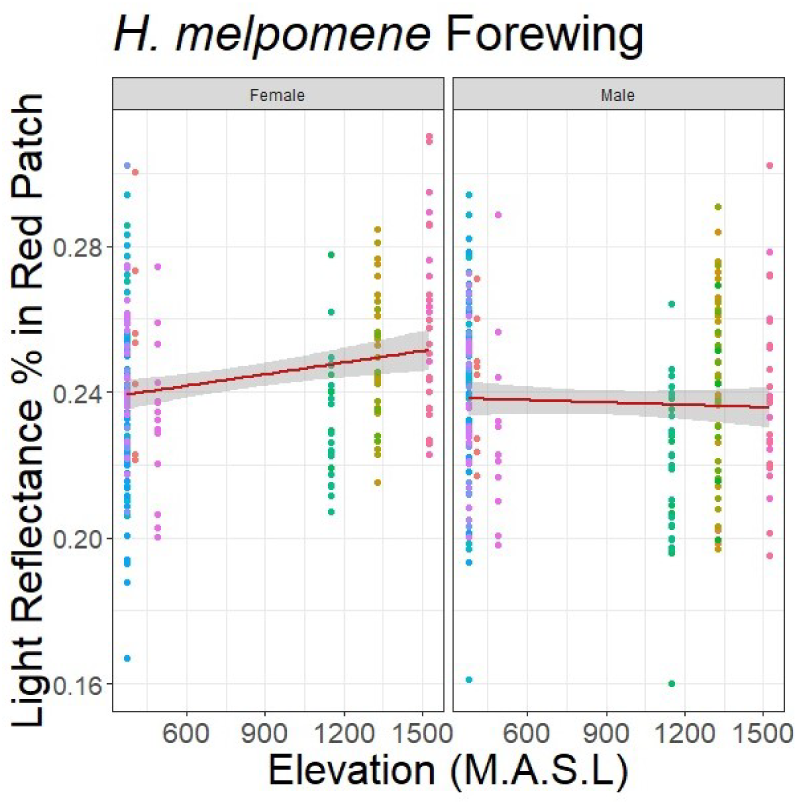
The relationship between reflectance of the forewing red wing patch (summed RGB values from colour photographs as a percentage of white values) with altitude in *H. melpomene,* separated by sex. Points are coloured by mother ID (points of the same colour are siblings). Trend lines are based on the raw individual data (not the effect sizes from the full model of all effects).

**Figure S3:**
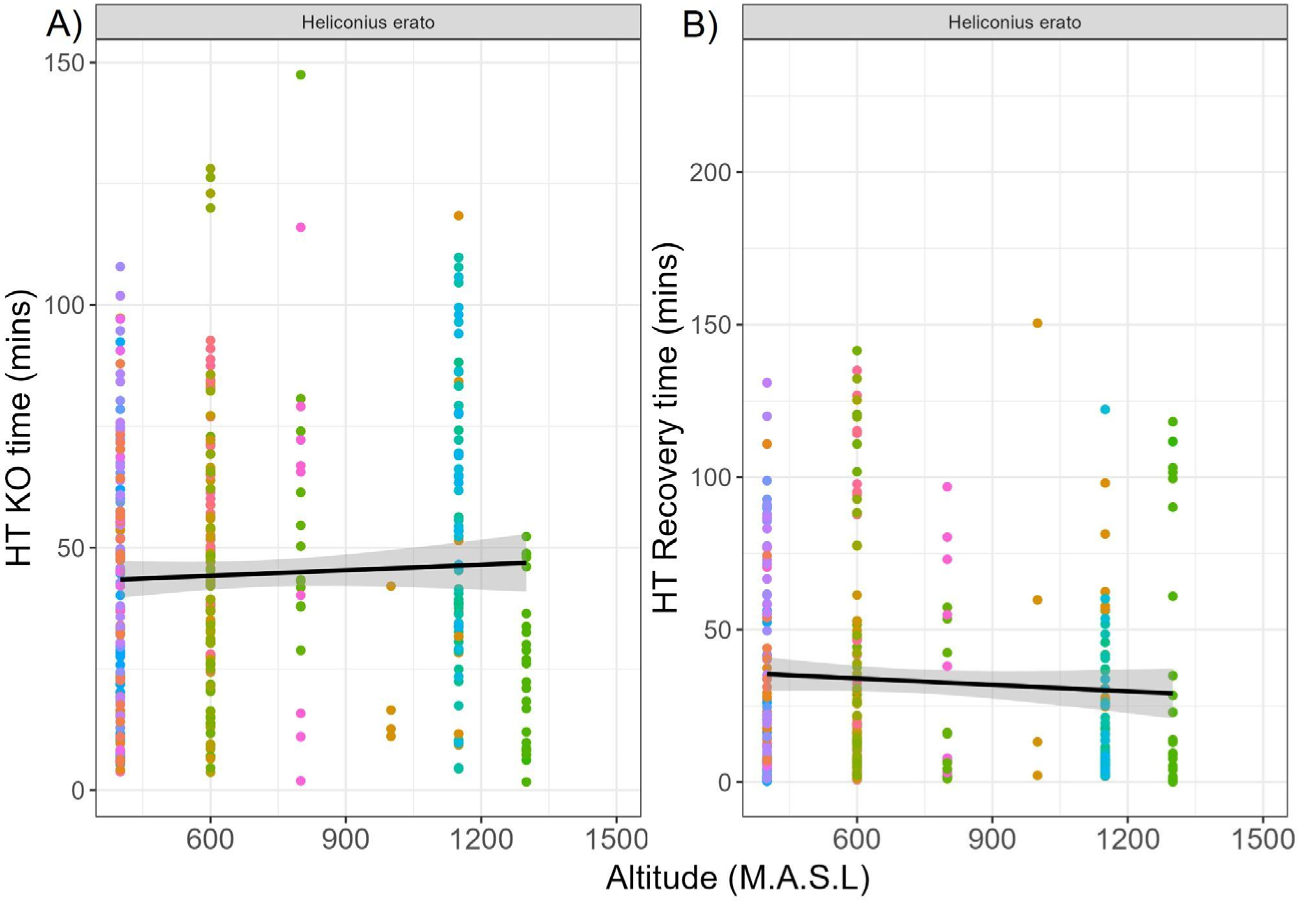
The relationships between heat Knock-out time (A) and recovery time from knock-out recovery (B) with altitude in *H. erato*. Longer knock-out times and shorter recovery times are both indicative of greater heat tolerance. Points are coloured by mother ID (points of the same colour are siblings). Trend lines are based on the raw individual data (not the effect sizes from the full model of all effects).

**Figure S4:**
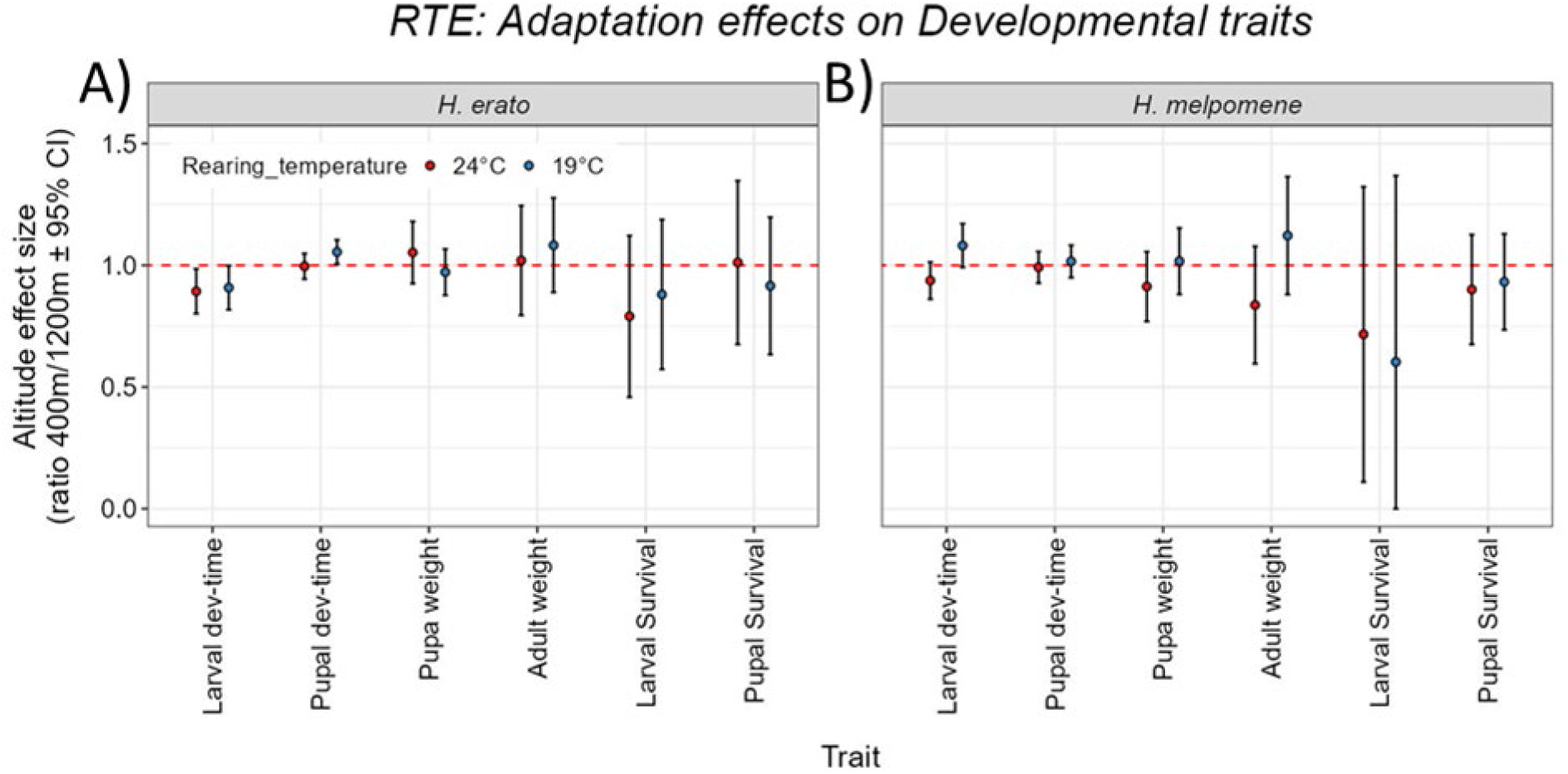
The effect of altitude in *H. erato* (A) and *H. melpomene* on their developmental traits as a ratio of 400m/1200m when reared at 24°C (red) and 19°C (blue)

**Figure S5:**
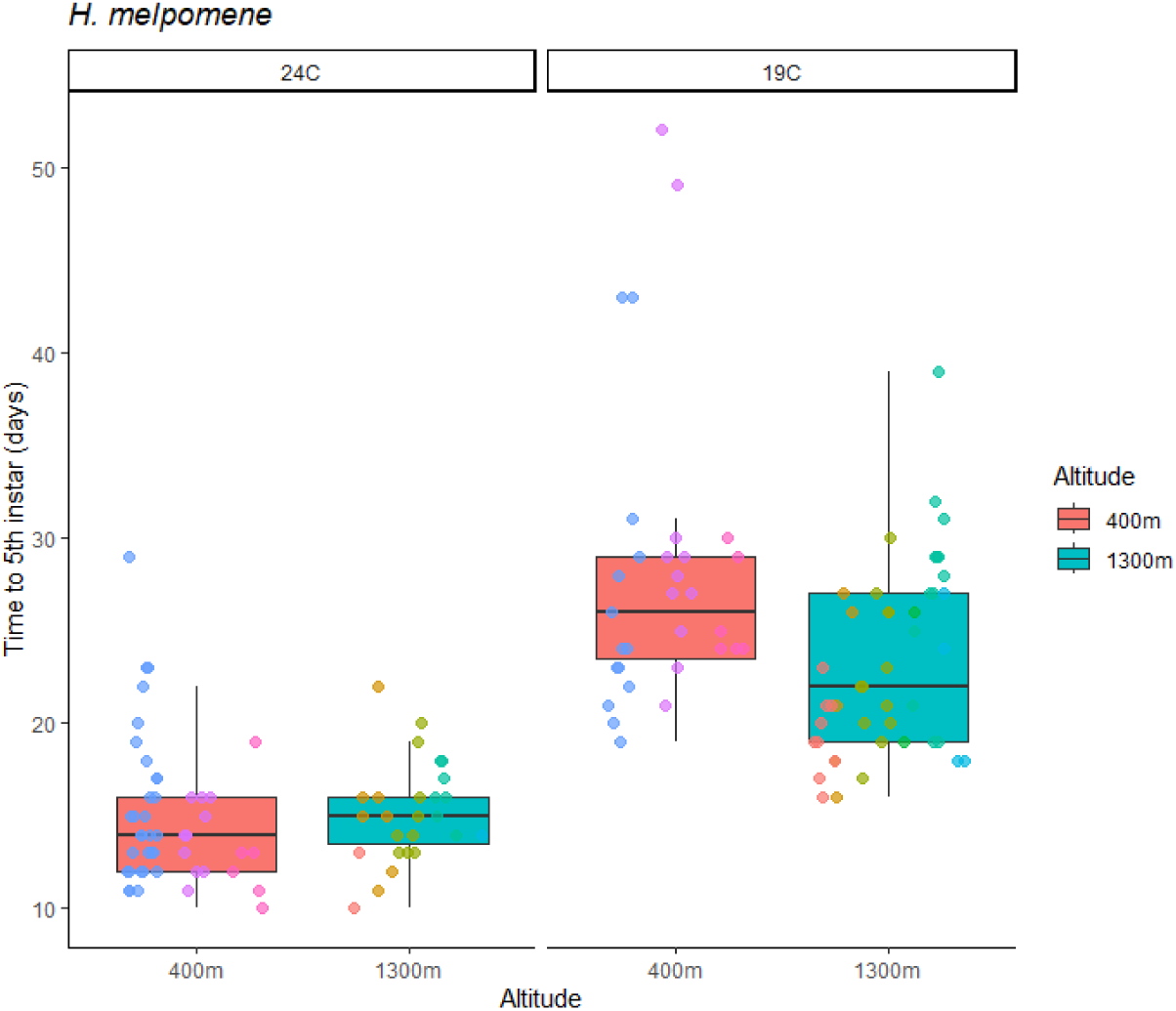
The effect of temperature and altitude on larval development time in *H. melpomene*. Boxes show the median and interquartile ranges for each population under each treatment, with overlain points for each individual. Boxes are coloured by population; points are coloured by mother ID (i.e. points of the same colour are siblings).

**Figure S6:**
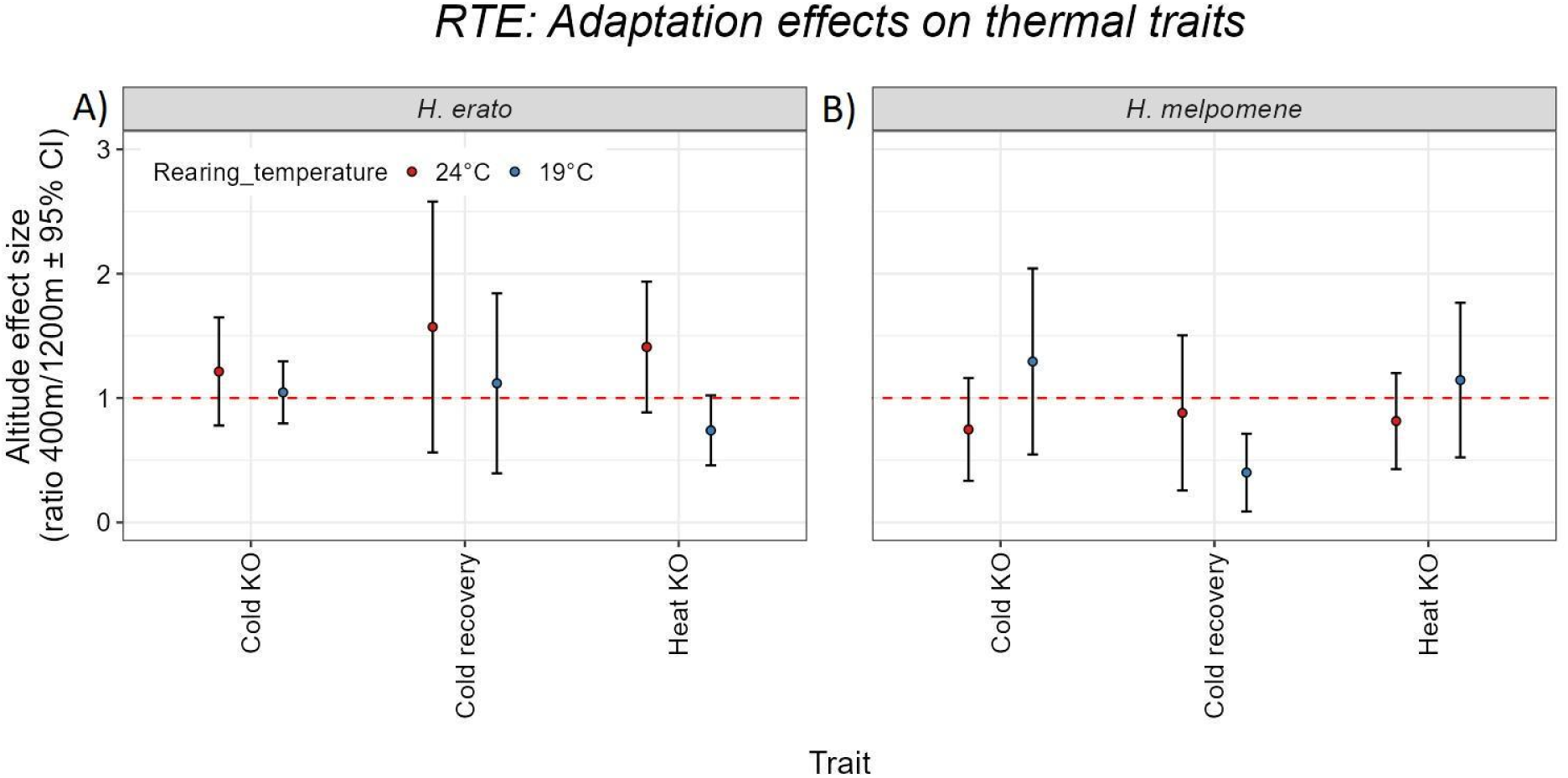
The effect of altitude in *H. erato* (A) and *H. melpomene* on their thermal tolerance traits as a ratio of 400m/1200m when reared at 24°C (red) and 19°C (blue).

**Figure S7:**
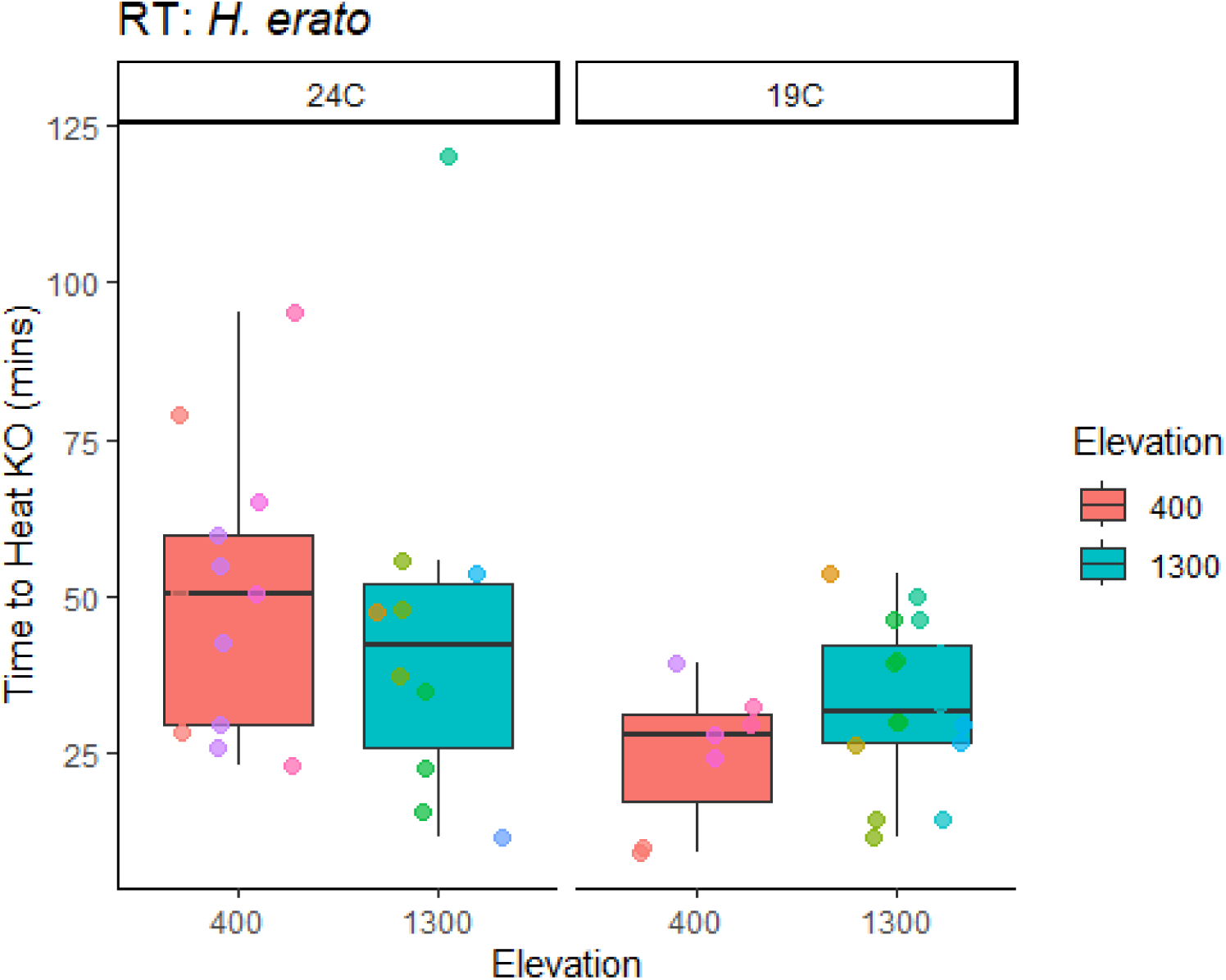
The effect of rearing temperature and altitude on time to heat knock-out at 39°C in *H. erato*. Boxes show the median and interquartile ranges for each population under each treatment, with overlain points for each individual. Boxes are coloured by population; points are coloured by mother ID (i.e. points of the same colour are siblings).

**Figure S8:**
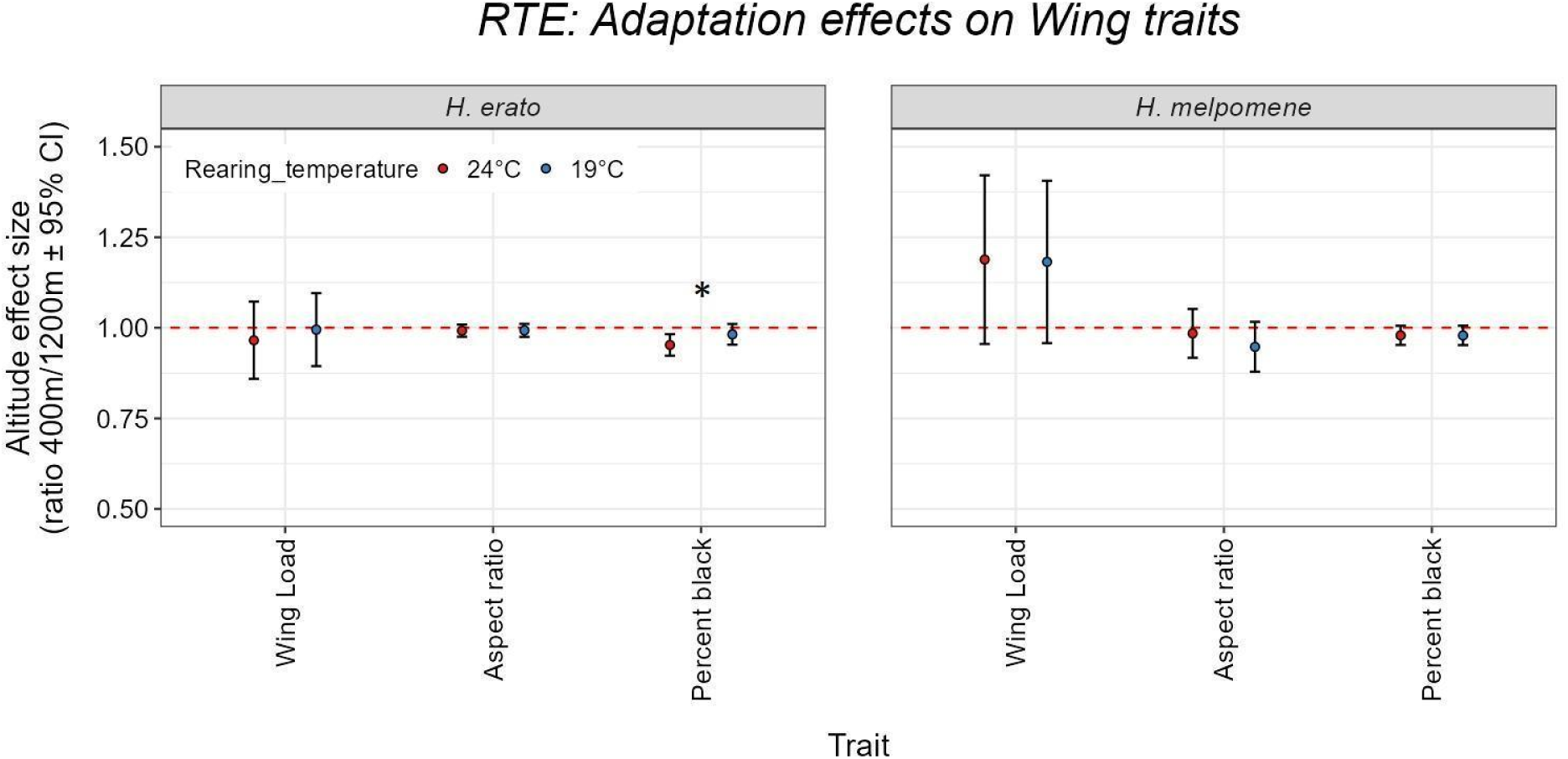
The effect of altitude in *H. erato* (A) and *H. melpomene* on their wing traits as a ratio of 400m/1200m when reared at 24°C (red) and 19°C (blue). Asterisk denotes that the statistical modelling supported an interaction between altitude and rearing temperature.

**Figure S9.**
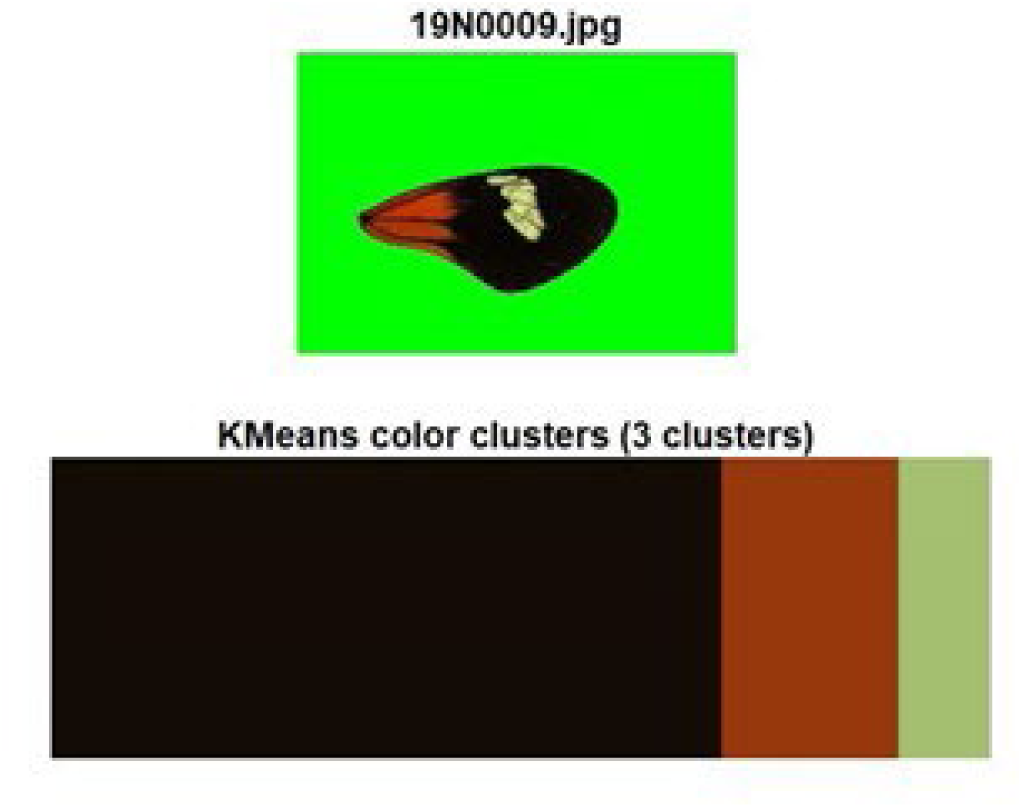
Example of a forewing image that was used for colour and shape analysis and the 3 colour bins that were used to quantify the % of the wing that was black and the reflectance of the “red” region (here appearing slightly orange/brown).

**Figure S10:**
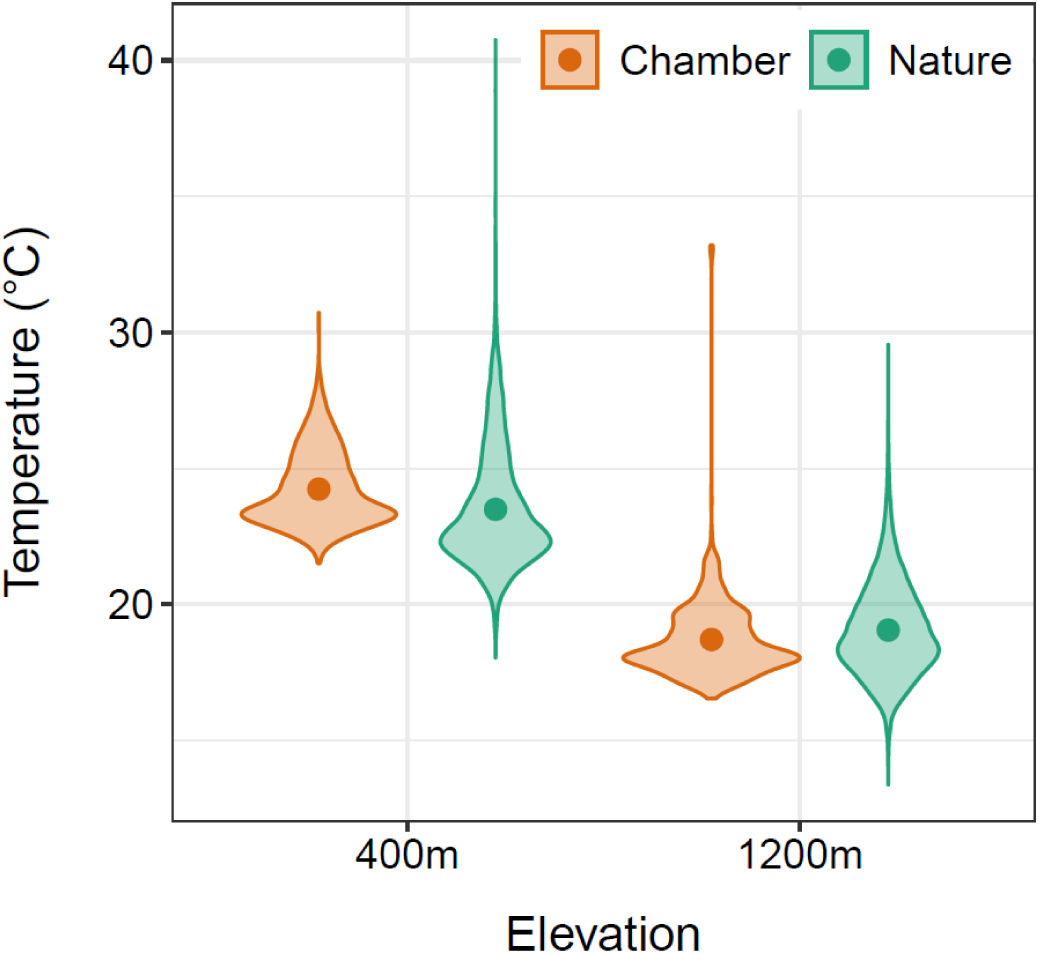
Thermal conditions in nature and in experimental chambers used for the reciprocal transplant experiment. Violin plots show temperature distributions (shape) and mean values (darker dots) at the elevations where butterflies (mothers) were originally collected and in the chambers simulating these environments. Both the average temperatures (19 °C at 400 m and 24 °C at 1200 m) and their variation were closely replicated in the experimental setup. Natural temperature data were recorded hourly over a full year at multiple forest understory sites, as reported by Montejo-Kovacevich et al.^28^

**Table S1:** Models used to test the effect of elevation in the GCE, per species, per trait. Delta AICc and AICc weights are given for the strongest model (the model with the lowest AICc, or that with the fewest explanatory variables within 2 AICc units of the model with the lowest AICc), and the model used to calculate the reported effect sizes. All fixed effects that were considered, and the sample numbers, are also reported.

| species | Trait | strongest model (with fewest explanatory factors) | $\Delta AICc$ | wt AICc | all fixed effects tested | Sample sizes | Effect size model | $\Delta AICc$ | wt AICc |
| --- | --- | --- | --- | --- | --- | --- | --- | --- | --- |
| *H. erato* | Laval development time | null | 0 | 0.66 | elevation | 38 mothers, 775 inds | ele | 1.35 | 0.34 |
| *H. melpomene* | Laval development time | null | 0 | 0.68 | elevation | 25 mothers, 450 inds | ele | 1.54 | 0.32 |
| *H. erato* | pupal development time | null | 0 | 0.7 | elevation | 38 mothers 558 inds | ele | 1.74 | 0.3 |
| *H. melpomene* | pupal development time | null | 0 | 0.7 | elevation | 22 mothers 405 inds | ele | 1.71 | 0.3 |
| *H. erato* | pupa weight | dev_time | 0 | 0.47 | elevation, development time | 40 mothers, 753 inds | ele + dev_time | 1.58 | 0.21 |
| *H. melpomene* | pupa weight | dev_time | 0 | 0.67 | elevation, development time | 23 mothers, 444 inds | ele + dev_time | 2.02 | 0.24 |
| *H. erato* | adult weight | dev_time | 0 | 0.56 | elevation, development time, sex | 45 mothers, 327 inds | ele + dev_time | 1.97 | 0.21 |
| *H. melpomene* | adult weight | dev_time | 0 | 0.3 | elevation, development time, sex | 29 mothers, 293, inds | ele + dev_time | 1.79 | 0.12 |
| *H. erato* | survival to pupation | null | 0 | 0.73 | elevation | 42 mothers, 1549 inds | ele | 2 | 0.27 |
| *H. melpomene* | survival to pupation | null | 0 | 0.7 | elevation | 26 mothers, 855 inds | ele | 1.74 | 0.3 |
| *H. erato* | survival post pupation | null | 0 | 0.73 | elevation | 40 mothers, 802 inds | ele | 2.01 | 0.27 |
| *H. melpomene* | survival post pupation | null | 0 | 0.6 | elevation | 24 mothers, 495 inds | ele | 0.77 | 0.4 |
| *H. erato* | wing load | null | 0 | 0.47 | elevation, development time, sex | 44 mothers, 319 inds | ele | 1.77 | 0.19 |
| *H. melpomene* | wing load | null | 0 | 0.4 | elevation, development time, sex | 29 mothers, 290 indis | ele | 1.48 | 0.19 |
| *H. erato* | black perc | SEX + ELE + W_Area | 0.5 | 0.14 | Elevation, Sex, Wing area | 36 mothers, 315 inds | same | same | same |
| *H. melpomene* | black perc | ELE + W_Area | 1.63 | 0.1 | Elevation, Sex, Wing area | 23 mothers, 333 inds | same | same | same |
| *H. erato* | red reflectance | SEX | 0.53 | 0.34 | Elevation, Sex | 33 mothers, 416 inds | same | same | same |
| *H. melpomene* | red reflectance | ELE*SEX | 0 | 0.75 | Elevation, Sex | 22 mothers, 431 inds | same | same | same |
| *H. erato* | Cold KO | AR + ELE + P_WEI + B.perc | 0 | 0.14 | Elevation, Observer, Black wing percentage, pupa weight, sex, wing aspect ratio | 37 mothers, 436 inds | same | same | same |
| *H. melpomene* | Cold KO | AR + ELE + HT_observer + P_WEI + B.perc | 0 | 0.07 | Elevation, Observer, Black wing percentage, pupa weight, sex, wing aspect ratio | 23 mothers, 374 inds | same | same | same |
| *H. erato* | Cold Recov | B.perc | 0 | 0.6 | Elevation, Observer, Black wing percentage, pupa weight, sex, wing aspect ratio | 37 mothers, 436 inds | same | same | same |
| *H. melpomene* | Cold Recov | AR + ELE + HT_observer + P_WEI + B.perc | 0 | 0.6 | Elevation, Observer, Black wing percentage, pupa weight, sex, wing aspect ratio | 23 mothers, 374 inds | same | same | same |
| *H. erato* | Heat KO | AR + ELE + SEX + P_WEI + B.perc + ELE:B.perc + AR:B.perc | 1.18 | 0.08 | Elevation, Observer, Black wing percentage, pupa weight, sex, wing aspect ratio | 31 mothers, 297 inds | same | same | same |
| *H. melpomene* | Heat KO | AR + ELE + HT_observer + P_WEI + B.perc | 0.37 | 0.07 | Elevation, Observer, Black wing percentage, pupa weight, sex, wing aspect ratio | 20 mothers, 239 inds | same | same | same |
| *H. erato* | Heat Recov | AR + ELE + HT_observer + P_WEI + B.perc + ELE:P_WEI | 0 | 0.09 | Elevation, Observer, Black wing percentage, pupa weight, sex, wing aspect ratio | 31 mothers, 297 inds | same | same | same |
| *H. melpomene* | Heat Recov | AR + ELE + HT_observer + P_WEI + B.perc | 0.81 | 0.04 | Elevation, Observer, Black wing percentage, pupa weight, sex, wing aspect ratio | 20 mothers, 239 inds | same | same | same |

**Table S2:** Heat tolerance models for Time to heat knock-out results with non-knockouts included in analysis.

| Experiment | species | Trait | strongest model<br>(with fewest<br>explanatory factors) | $\Delta AICc$ | wt AICc | all fixed effects | n numbers |
| --- | --- | --- | --- | --- | --- | --- | --- |
| CGE | *H. erato* | HT_KO | AR + ELE + SEX + P_WEI<br>+ B.perc + ELE:B.perc | 0 | 0.15 | Elevation, Observer, Black wing percentage,<br>pupa weight, sex, wing aspect ratio | 31 mothers,<br>367 inds |
| CGE | *H. melpomene* | HT_KO | AR + ELE + HT_observer<br>+ P_WEI + B.perc +<br>ELE:SEX | 0 | 0.06 | Elevation, Observer, Black wing percentage,<br>pupa weight, sex, wing aspect ratio | 20 mothers,<br>275 inds |
| RTE | *H. erato* | HT_KO | TEM*ELE + Observer | 0 | 0.93 | Temperature, Elevation, Observer | 12 mothers,<br>46 inds |

**Table S3:** Models used to test the effects of temperature and elevation in the RTE, per species, per trait. Delta AICc and AICc weights are given for the strongest model (the model with the lowest AICc, or that with the fewest explanatory variables within 2 AICc units of the model with the lowest AICc), and the model used to calculate the reported effect sizes. All fixed effects that were considered, and the sample numbers, are also reported.

| species | Trait | strongest model<br>(with fewest<br>explanatory factors) | $\Delta AICc$ | wt<br>AICc | all fixed effects | n numbers | Effect size model | $\Delta AICc$ | wt<br>AICc |
| --- | --- | --- | --- | --- | --- | --- | --- | --- | --- |
| *H. erato* | larval<br>development<br>time | TEM | 1.44 | 0.26 | Temperature,<br>Elevation, | 16 mothers,<br>199 inds | TEM*ELE | 1.92 | 0.2 |
| *H. melpomene* | larval<br>development<br>time | TEM*ELE | 0 | 0.78 | Temperature,<br>Elevation, | 8 mothers,<br>79 inds | same | same | same |
| *H. erato* | Pupal development time | TEM | 1.77 | 0.15 | Temperature, Elevation, Sex | 15 mothers, 143 inds | TEM*ELE | 0 | 0.35 |
| *H. melpomene* | Pupal development time | TEM | 0 | 0.46 | Temperature, Elevation, Sex | 8 mothers, 68 inds | TEM*ELE | 3.99 | 0.06 |
| *H. erato* | pupal weight | TEM*Dev_time | 0 | 0.46 | Temperature, Elevation, developmental time | 16 mothers, 196 inds | TEM, ELE +DEV_TIME+ TEM:ELE+ TEM:DEV_TIME | 1.82 | 0.18 |
| *H. melpomene* | pupal weight | TEM, ELE, DEV_TIME, TEM:DEV_TIME, ELE:DEV_TIME | 0 | 0.56 | Temperature, Elevation, developmental time | 8 mothers, 75 inds | TEM+ELE+ DEV_TIME+ TEM:ELE+ TEM:DEV_TIME+ ELE:DEV_TIME | 1.34 | 0.29 |
| *H. erato* | adult weight | TEM | 1.61 | 0.11 | Temperature, Elevation, developmental time, Sex | 15 mothers, 131 inds | TEM, ELE, DEV_TIME, TEM:ELE, TEM:DEV_TIME | 3.41 | 0.05 |
| *H. melpomene* | adult weight | ELE:DEV_TIME +ELE +DEV_TIME | 0.48 | 0.21 | Temperature, Elevation, developmental time, sex | 8 mothers, 58 inds | TEM+ELE+ DEV_TIME+ TEM:ELE+ TEM:DEV_TIME+ ELE:DEV_TIME | 0 | 0.27 |
| *H. erato* | larval survival | null | 0 | 0.46 |  | 16 mothers, 297 inds | TEM*ELE | 4.25 | 0.06 |
| *H. melpomene* | larval survival | null | 0 | 0.37 |  | 9 mothers, 155 inds | TEM*ELE | 3.2 | 0.08 |
| *H. erato* | pupal survival | null | 0 | 0.54 |  | 16 mothers, 200 inds | TEM*ELE | 5.66 | 0.03 |
| *H. melpomene* | pupal survival | null | 0 | 0.38 |  | 9 mothers, 116 inds | TEM*ELE | 3.72 | 0.06 |
| *H. erato* | Wing load<br>mg/mm | Dev_time | 0 | 0.19 | Temperature,<br>elevation, sex,<br>development time | 14 mothers,<br>127 inds | TEM:ELE +Dev_time + ELE<br>+ TEM | 4.68 | 0.02 |
| *H. melpomene* | Wing load<br>mg/mm | ELE:Dev_time +<br>ELE:SEX + TEM +ELE +<br>SEX +Dev_time | 0 | 0.13 | Temperature,<br>elevation, sex,<br>development time | 8 mothers,<br>87 inds | ELE:Dev_time + ELE:SEX +<br>ELE:TEM + TEM +ELE +<br>SEX +Dev_time | 1.58 | 0.06 |
| *H. erato* | black perc | TEM:ELE + TEM:SEX +<br>TEM + ELE + SEX +<br>W_Area | 0 | 0.19 | Temperature,<br>Elevation, Sex, wing<br>area | 13 mothers,<br>92 inds | same | same | same |
| *H. melpomene* | black perc | TEM:W_Area +<br>W_area:SEX + TEM +<br>ELE + SEX + W_area | 0 | 0.14 | Temperature,<br>Elevation, Sex, wing<br>area | 8 mothers,<br>75 inds | same | same | same |
| *H. erato* | Aspect ratio | null | 1.38 | 0.1 | Temperature,<br>elevation, sex, adult<br>weight | 14 mothers,<br>127 inds | TEM*ELE | 6.06 | 0.01 |
| *H. melpomene* | Aspect ratio | ELE:SEX + SEX:A_Wei +<br>TEM +ELE + SEX +<br>A_Wei | 0 | 0.15 | Temperature,<br>elevation, sex, adult<br>weight | 8 mothers,<br>87 inds | ELE:SEX + SEX:A_Wei +<br>ELE:TEM+ TEM +ELE +<br>SEX + A_Wei | 9.61 | 0 |
| *H. erato* | CT_KO | TEM | 0 | 0.51 | Temperature,<br>Elevation, Observer | 12 mothers,<br>35 inds | TEM*ELE | 6.23 | 0.02 |
| *H. melpomene* | CT_KO | null | 0 | 0.5 | Temperature,<br>Elevation, Observer | 6 mothers,<br>23 inds | TEM*ELE | 4.19 | 0.03 |
| *H. erato* | CT_Recov | null | 0 | 0.47 | Temperature,<br>Elevation, Observer | 12 mothers,<br>35 inds | TEM*ELE | 3.95 | 0.07 |
| *H. melpomene* | CT_Recov | NULL | 0 | 0.28 | Temperature,<br>Elevation, Observer | 6 mothers,<br>23 inds | TEM*ELE | 6.12 | 0.02 |
| *H. erato* | HT_KO | TEM*ELE + Observer | 0 | 0.67 | Temperature,<br>Elevation, Observer | 12 mothers,<br>45 inds | same | same | same |
| *H. melpomene* | HT_KO | null | 0 | 0.35 | Temperature,<br>Elevation, Observer | 6 mothers,<br>35 inds | TEM*ELE | same | same |

## References

1. Humboldt, A. von & Bonpland, A. Essay on the Geography of Plants. (1807).

2. Hodkinson, I. D. Terrestrial insects along elevation gradients: species and community responses to altitude. Biol. Rev. 80, 489–513 (2005).

3. Janzen, D. H. Why Mountain Passes are Higher in the Tropics. Am. Nat. 101, 233–249 (1967).

4. Ghalambor, C. K. Are mountain passes higher in the tropics? janzen’s hypothesis revisited. Integr. Comp. Biol. 46, 5–17 (2006).

5. Slade, E. M. & Ong, X. R. The future of tropical insect diversity: strategies to fill data and knowledge gaps. Curr. Opin. Insect Sci. 58, 101063 (2023).

6. Körner, C. The use of ‘altitude’ in ecological research. Trends Ecol. Evol. 22, 569–574 (2007).

7. Sullivan, J. B. & Miller, W. E. Intraspecific body size variation in macrolepidoptera as related to altitude of capture site and seasonal generation. J. Lepid. Soc. 61, 72–77 (2007).

8. Bergmann, C. Über Die Verhältnisse Der Wärmeökonomie Der Thiere Zu Ihrer Grösse. (1848).

9. Scharf, I., Sbilordo, S. H. & Martin, O. Y. Cold tolerance in flour beetle species differing in body size and selection temperature. Physiol. Entomol. 39, 80–87 (2014).

10. Brehm, G., Zeuss, D. & Colwell, R. K. Moth body size increases with elevation along a complete tropical elevational gradient for two hyperdiverse clades. Ecography (Cop.). 42, 632–642 (2019).

11. Nève, G. & Després, L. Cold adaptation across the elevation gradient in an alpine butterfly species complex. Ecol. Entomol. 45, 997–1003 (2020).

12. Dufour, P. C. et al. Divergent melanism strategies in Andean butterfly communities structure diversity patterns and climate responses. J. Biogeogr. 45, 2471–2482 (2018).

13. Bogert, C. M. Thermoregulation in Reptiles, A Factor in Evolution. Evolution (N. Y). 3, 195 (1949).

14. Clusella Trullas, S., van Wyk, J. H. & Spotila, J. R. Thermal melanism in ectotherms. J. Therm. Biol. 32, 235–245 (2007).

15. Fox, R. J., Donelson, J. M., Schunter, C., Ravasi, T. & Gaitán-Espitia, J. D. Beyond buying time: the role of plasticity in phenotypic adaptation to rapid environmental change. Philos. Trans. R. Soc. B Biol. Sci. 374, (2019).

16. Karl, I. et al. Altitudinal life-history variation and thermal adaptation in the copper butterfly Lycaena tityrus. Oikos 117, 778–788 (2008).

17. Kingsolver, J. G. & Buckley, L. B. Evolution of plasticity and adaptive responses to climate change along climate gradients. Proc. R. Soc. B Biol. Sci. 284, (2017).

18. Niitepõld, K. Genotype by temperature interactions in the metabolic rate of the Glanville fritillary butterfly. J. Exp. Biol. 213, 1042–1048 (2010).

19. Brakefield, P. M. & Kesbeke, F. Genotype-environment interactions for insect growth in constant and fluctuating temperature regimes. Proc. R. Soc. London. Ser. B Biol. Sci. 264, 717–723 (1997).

20. Gienapp, P., Teplitsky, C., Alho, J. S., Mills, J. A. & Merilä, J. Climate change and evolution: disentangling environmental and genetic responses. Mol. Ecol. 17, 167– 178 (2008).

21. Murren, C. J. et al. Constraints on the evolution of phenotypic plasticity: limits and costs of phenotype and plasticity. Hered. 2015 1154 115, 293–301 (2015).

22. Leroi, A. M., Bennett, A. F. & Lenski, R. E. Temperature acclimation and competitive fitness: an experimental test of the beneficial acclimation assumption. Proc. Natl. Acad. Sci. U. S. A. 91, 1917 (1994).

23. Wilson, R. S. & Franklin, C. E. Testing the beneficial acclimation hypothesis. Trends Ecol. Evol. 17, 66–70 (2002).

24. Sejerkilde, M., Sørensen, J. G. & Loeschcke, V. Effects of cold- and heat hardening on thermal resistance in Drosophila melanogaster. J. Insect Physiol. 49, 719–726 (2003).

25. Fischer, K. et al. Environmental Effects on Temperature Stress Resistance in the Tropical Butterfly Bicyclus Anynana. PLoS One 5, e15284 (2010).

26. Franke, K. et al. Effects of adult temperature on gene expression in a butterfly: Identifying pathways associated with thermal acclimation. BMC Evol. Biol. 19, 1–16 (2019).

27. Rosser, N., Phillimore, A. B., Huertas, B., Willmott, K. R. & Mallet, J. Testing historical explanations for gradients in species richness in heliconiine butterflies of tropical America. Biol. J. Linn. Soc. 105, 479–497 (2012).

28. Montejo-Kovacevich, G. et al. Microclimate buffering and thermal tolerance across elevations in a tropical butterfly. J. Exp. Biol. 223, (2020).

29. Montejo-Kovacevich, G. et al. Repeated genetic adaptation to altitude in two tropical butterflies. Nat. Commun. 2022 131 13, 1–16 (2022).

30. Montejo-Kovacevich, G. et al. Genomics of altitude-associated wing shape in two tropical butterflies. Mol. Ecol. 30, 6387–6402 (2021).

31. Montejo-Kovacevich, G. et al. Altitude and life-history shape the evolution of *Heliconius* wings. Evolution (N. Y). 73, 2436–2450 (2019).

32. Falconer, D. S. & Mackay, T. F.. *Introduction to Quantitative Genetics*. (Pearson, 1996).

33. Walters, J. R., Corbins, C., Hardcastle, T. J. & Jiggins, C. D. Evaluating female remating rates in light of spermatophore degradation in Heliconius butterflies: pupal-mating monandry versus adult-mating polyandry. Ecol. Entomol. 37, 257–268 (2012).

34. Dasmahapatra, K. K. et al. Butterfly genome reveals promiscuous exchange of mimicry adaptations among species. Nature 487, 94–98 (2012).

35. Nadeau, N. J. Genes controlling mimetic colour pattern variation in butterflies. Curr. Opin. Insect Sci. 17, 24–31 (2016).

36. Bainbridge, H. E. et al. Limited genetic parallels underlie convergent evolution of quantitative pattern variation in mimetic butterflies. J. Evol. Biol. 33, 1516–1529 (2020).

37. Günter, F. et al. Latitudinal and altitudinal variation in ecologically important traits in a widespread butterfly. Biol. J. Linn. Soc. 128, 742–755 (2019).

38. Barton, M. & Porter, W. Behavioural thermoregulation and the relative roles of convection and radiation in a basking butterfly. J. Therm. Biol. 41, 65–71 (2014).

39. Kuyucu, A. C., Sahin, M. K. & Caglar, S. S. The relation between melanism and thermal biology in a colour polymorphic bush cricket, Isophya rizeensis. J. Therm. Biol. 71, 212–220 (2018).

40. Klein, A. L. & de Araújo, A. M. Sexual Size Dimorphism in the Color Pattern Elements of Two Mimetic Heliconius Butterflies. Neotrop. Entomol. 42, 600–606 (2013).

41. Jiggins, C. D. *The Ecology and Evolution of Heliconius Butterflies*. (OUP Oxford, Oxford, 2017). doi:10.1093/acprof:oso/9780199566570.001.0001.

42. Kristensen, T. N. et al. Costs and benefits of cold acclimation in field-released Drosophila. Proc. Natl. Acad. Sci. U. S. A. 105, 216–221 (2008).

43. Karl, I. et al. Interactive effects of acclimation temperature and short-term stress exposure on resistance traits in the butterfly Bicyclus anynana. Physiol. Entomol. 39, 222–228 (2014).

44. Reid, K., Bell, M. A. & Veeramah, K. R. Threespine Stickleback: A Model System for Evolutionary Genomics. Annu. Rev. Genomics Hum. Genet. 22, 357–383 (2021).

45. Leingärtner, A., Krauss, J. & Steffan-Dewenter, I. Species richness and trait composition of butterfly assemblages change along an altitudinal gradient. Oecologia 175, 613–623 (2014).

46. Sinclair, B. J., Williams, C. M. & Terblanche, J. S. Variation in Thermal Performance among Insect Populations. Physiol. Biochem. Zool. 85, 594–606 (2012).

47. Horne, C. R., Hirst, A. G. & Atkinson, D. Insect temperature–body size trends common to laboratory, latitudinal and seasonal gradients are not found across altitudes. Funct. Ecol. 32, 948–957 (2018).

48. Ashton, K. G. Patterns of within-species body size variation of birds: strong evidence for Bergmann’s rule. Glob. Ecol. Biogeogr. 11, 505–523 (2002).

49. Svensson, E. I., Gómez-Llano, M. & Waller, J. T. Out of the tropics: Macroevolutionary size trends in an old insect order are shaped by temperature and predators. J. Biogeogr. 50, 489–502 (2023).

50. Mousseau, T. A. Ectotherms Follow the Converse to Bergmann’s Rule. Evolution (N. Y). 51, 630 (1997).

51. Atkinson, D. Temperature and Organism Size—A Biological Law for Ectotherms? Adv. Ecol. Res. 25, 1–58 (1994).

52. Kingsolver, J. G. & Huey, R. B. Size, temperature, and fitness: Three rules. Evol. Ecol. Res. 10, 251–268 (2008).

53. Teplitsky, C. & Millien, V. Climate warming and Bergmann’s rule through time: is there any evidence? Evol. Appl. 7, 156 (2014).

54. Kemp, D. J. & Krockenberger, A. K. Behavioural thermoregulation in butterflies: the interacting effects of body size and basking posture in Hypolimnas bolina (L.) (Lepidoptera : Nymphalidae). Aust. J. Zool. 52, 229–239 (2004).

55. Klockmann, M., Günter, F. & Fischer, K. Heat resistance throughout ontogeny: body size constrains thermal tolerance. Glob. Chang. Biol. 23, 686–696 (2017).

56. Mallet, J. et al. Estimates of selection and gene flow from measures of cline width and linkage disequilibrium in heliconius hybrid zones. Genetics 124, 921–936 (1990).

57. Semper, C. The Natural Conditions of Existence as They Affect Animal Life. (1883).

58. Weller, H. I. & Westneat, M. W. Quantitative color profiling of digital images with earth mover’s distance using the R package colordistance. PeerJ 7, (2019).

59. Burnham, K. P. & Anderson, D. R. Model Selection and Multimodel Inference. (2004).

60. Bates, D., Mächler, M., Bolker, B. M. & Walker, S. C. Fitting Linear Mixed-Effects Models Using lme4. J. Stat. Softw. 67, 1–48 (2015).

61. Mazerolle, M. J. CRAN - Package AICcmodavg. at https://cran.r-project.org/web/packages/AICcmodavg/index.html (2023).

62. Nakagawa, S. & Schielzeth, H. Repeatability for Gaussian and non-Gaussian data: a practical guide for biologists. Biol. Rev. Camb. Philos. Soc. 85, 935–956 (2010).

63. Stoffel, M. A., Nakagawa, S. & Schielzeth, H. rptR: repeatability estimation and variance decomposition by generalized linear mixed-effects models. Methods Ecol. Evol. 8, 1639–1644 (2017).

64. Searle, S. R., Speed, F. M. & Milliken, G. A. Estimated Marginal Means, aka Least-Squares Means [R package emmeans version 1.8.9]. Am. Stat. 34, 216–221 (2023).

65. Wickham, H. ggpolt2 Elegant Graphics for Data Analysis. Use R! Ser. 211 (2016).

66. Massicotte, P. & South, A. rnaturalearth: World Map Data from Natural Earth. at https://docs.ropensci.org/rnaturalearth/,%0Ahttps://github.com/ropensci/rnaturalearth (2023).

67. Hijmans, R. J. Geographic Data Analysis and Modeling [R package raster version 3.6-26]. (2023).

68. Pebesma, E. & Bivand, R. Spatial Data Science: With Applications in R. Spat. Data Sci. With Appl. R 1–300 (2023) doi:10.1201/9780429459016.

69. Pebesma, E. Simple features for R: Standardized support for spatial vector data. R J. 10, 439–446 (2018).

70. Slowikowski, K. Automatically Position Non-Overlapping Text Labels with ‘ggplot2’ [R package ggrepel version 0.9.4]. (2023).

